# Single-cell nascent RNA sequencing using click-chemistry unveils coordinated transcription

**DOI:** 10.1101/2023.09.15.558015

**Authors:** Dig B. Mahat, Nathaniel D. Tippens, Jorge D. Martin-Rufino, Sean K. Waterton, Jiayu Fu, Sarah E. Blatt, Phillip A. Sharp

## Abstract

Transcription is the primary regulatory step in gene expression. Divergent transcription initiation from promoters and enhancers produces stable RNAs from genes and unstable RNAs from enhancers^1–5^. Nascent RNA capture and sequencing assays simultaneously measure gene and enhancer activity in cell populations^6–9^. However, fundamental questions in the temporal regulation of transcription and enhancer-gene synchrony remain unanswered primarily due to the absence of a single-cell perspective on active transcription. In this study, we present scGRO-seq - a novel single-cell nascent RNA sequencing assay using click-chemistry - and unveil the coordinated transcription throughout the genome. scGRO-seq demonstrates the episodic nature of transcription, and estimates burst size and frequency by directly quantifying transcribing RNA polymerases in individual cells. It reveals the co-transcription of functionally related genes and leverages the replication-dependent non-polyadenylated histone genes transcription to elucidate cell-cycle dynamics. The single-nucleotide spatial and temporal resolution of scGRO-seq identifies networks of enhancers and genes and indicates that the bursting of transcription at super-enhancers precedes the burst from associated genes. By imparting insights into the dynamic nature of transcription and the origin and propagation of transcription signals, scGRO-seq demonstrates its unique ability to investigate the mechanisms of transcription regulation and the role of enhancers in gene expression.

## Main

Transcription is a discontinuous process characterized by short bursts and long inter-burst silent periods^10–14^. Decoding the origin and circuits of burst signals is critical in understanding the mechanisms of transcription regulation during the cell cycle, development, and diseases. Core promoter elements, transcription factors, and enhancers are implicated in regulating burst kinetics, but their precise role in determining the overall transcription output remains unsettled^15–18^. Whether the widely accepted view of stochastic transcription of individual genes conceals synchronous transcription of functionally related genes and coordination between enhancer-gene pairs holds broad significance in understanding gene regulation. From a clinical perspective, assessing the contribution of enhancers in regulating protein-coding genes can unlock a largely unexplored genomic landscape for therapeutics.

Active enhancers are occupied by transcription factors and RNA polymerase, similar to the gene promoters they regulate, resulting in the synthesis of non-coding, non-polyadenylated, and unstable RNA^3,19,20^. They are highly specific to cell types and states^21^, exerting cis-regulatory effects over long genomic distances^22^. Genome-wide association studies further underscore the role of enhancers in gene regulation, showing that over 90% of genomic loci associated with traits and diseases are found in non-coding regions with many overlapping enhancers^23^. However, linking enhancers harboring causal variants to genes remains challenging. Although low throughput, genome editing tools can potentially map enhancer-gene pairs, but the pleiotropic nature^17^ and weak effect of individual enhancers hinder their utility.

Existing genomic tools that probe the coding and non-coding genome without perturbation by assessing chromatin conformation, histone modifications, and chromatin accessibility shed light on the molecular events leading up to the enhancer-mediated gene activation. However, these tools do not fully confirm the actual activation^24^. Despite having similar chromatin features, the distinguishing feature of an active enhancer from its inactive counterpart is its transcription^25^. Nascent RNA sequencing assays, such as global run-on and sequencing (GRO-seq)^8^ and precision run-on and sequencing (PRO-seq)^9^, enable the simultaneous quantification of transcription in genes and enhancers. However, these bulk cell assays average the discontinuous transcription from individual cells, making it challenging to decipher transcription dynamics and assign enhancer-gene relationships.

Here, we present a novel single-cell nascent RNA sequencing (scGRO-seq) method using copper-catalyzed azide-alkyne cycloaddition (CuAAC or click-chemistry)^26^ to assess genome-wide nascent transcription in individual cells quantitatively. Our analyses of genes and enhancers across 2,635 individual mouse embryonic stem cells (mESCs) provide a comprehensive view of the dynamic nature of transcription. We leverage the elongating RNA polymerases as built-in clocks and measure the distance traveled from the transcription start sites (TSS) to estimate transcriptional burst kinetics. Using a class of cell-cycle phase-specific genes undetected by most single-cell methods, we quantify the dynamics of transcription during cell-cycle. We used the single-nucleotide temporal resolution of genome-wide transcription in individual cells to reveal the co-transcribed gene-gene and enhancer-gene networks that are turned on within a few minutes of each other. Using a set of validated enhancer-gene pairs, we show preliminary evidence for the transcription initiation at enhancers before the transcription activation at the associated genes. Overall, scGRO-seq bridges a critical gap in the study of temporal control of transcription and the functional association of enhancers and genes, and these insights will shed light on the gene regulatory mechanisms in essential cellular processes and disease.

### Development of scGRO-seq

The primary challenge in capturing and sequencing nascent RNA from individual cells is attaching unique single-cell tags onto nascent RNA. Existing nascent RNA sequencing methods selectively capture tagged-nascent RNA from a cell population, making single-cell deconvolution impossible. In contrast, single-cell RNA sequencing (scRNA-seq) methods capture mRNA by annealing with the polyA tail and attach single-cell barcode sequences by reverse transcription. Nascent RNA lacks a terminal polyA tract or other consensus sequence and must be selectively labeled and enriched from abundant total cellular RNA.

We designed a novel strategy to selectively label nascent RNA by nuclear run-on reaction in the presence of modified nucleotide triphosphates (NTPs) compatible with CuAAC conjugation. CuAAC is highly efficient, extremely selective, robust under diverse reaction conditions, enzyme-free and compatible with automation. First, we developed, optimized, and systematically characterized an Assay for Genome-wide Transcriptome using Click-chemistry (AGTuC) - a cell population-based nascent RNA sequencing method using 3’-(O-Propargyl)-NTPs in mESCs (Extended Data Fig. 1a-f, 2a-d). It takes ∼8 hours to prepare an AGTuC library. However, the high concentration of ionic detergent in AGTuC disrupts nuclear membranes during run-on reaction, making RNA from individual cells indistinguishable for single-cell barcoding. We therefore developed an iteration of AGTuC where nascent RNAs in individual nuclei are labeled with alkyne via run-on with 3’-(O-Propargyl)-NTPs but without disrupting the nuclear membrane (inAGTuC) (Extended Data Fig. 3a-j, 4a-h). We prepared inAGTuC libraries from ∼100K, ∼10K, and ∼1K nuclei (Extended Data Fig. 5a-d) and tested for correlation among itself and with PRO-seq, demonstrating the feasibility of profiling nascent RNA with small sample sizes. Based on the correlation slope, the inAGTuC library with as low as ∼1K nuclei showed similar efficiency as PRO-seq in detecting nascent transcriptomes. The higher efficiency, lower cost, shorter library preparation time, and lower sample input make AGTuC and inAGTuC viable alternatives to existing methods like PRO-seq. By enabling the compartmentalization of intact nuclei containing click-compatible nascent RNA and 5’- azide-single-cell-barcoded (5’-AzScBc) DNA molecules using fewer nuclei, inAGTuC laid the ground for single-cell nascent RNA sequencing.

Building on this foundation, we applied our newly developed chemistry to single cells. For congruence with the original nascent RNA sequencing method of GRO-seq, we named this single-cell version scGRO-seq (Fig. 1a). Intact nuclei containing nascent RNA labeled with propargyl, following a nuclear run-on reaction with 3’-(O-Propargyl)-NTPs are sorted individually into 96-well plates. Each well contains a small volume of 8 M urea, which lyses the nuclear membrane and denatures RNA polymerase, releasing propargyl-labeled nascent RNA. Adding CuAAC reagents covalently links propargyl-labeled nascent RNA to a unique 5’-AzScBc DNA molecule in each well. After CuAAC, single-cell-barcoded nascent RNAs from 96 wells are pooled, reverse transcribed in the presence of a template switching oligo (TSO), PCR amplified, and sequenced (Extended Data Fig. 6). Despite a span of more than three years between the generation of various scGRO-seq library replicates, the 36 batches that passed quality control showed strong correlation at the 96-well plate level (Extended Data Fig. 7a).

**Fig. 1.**
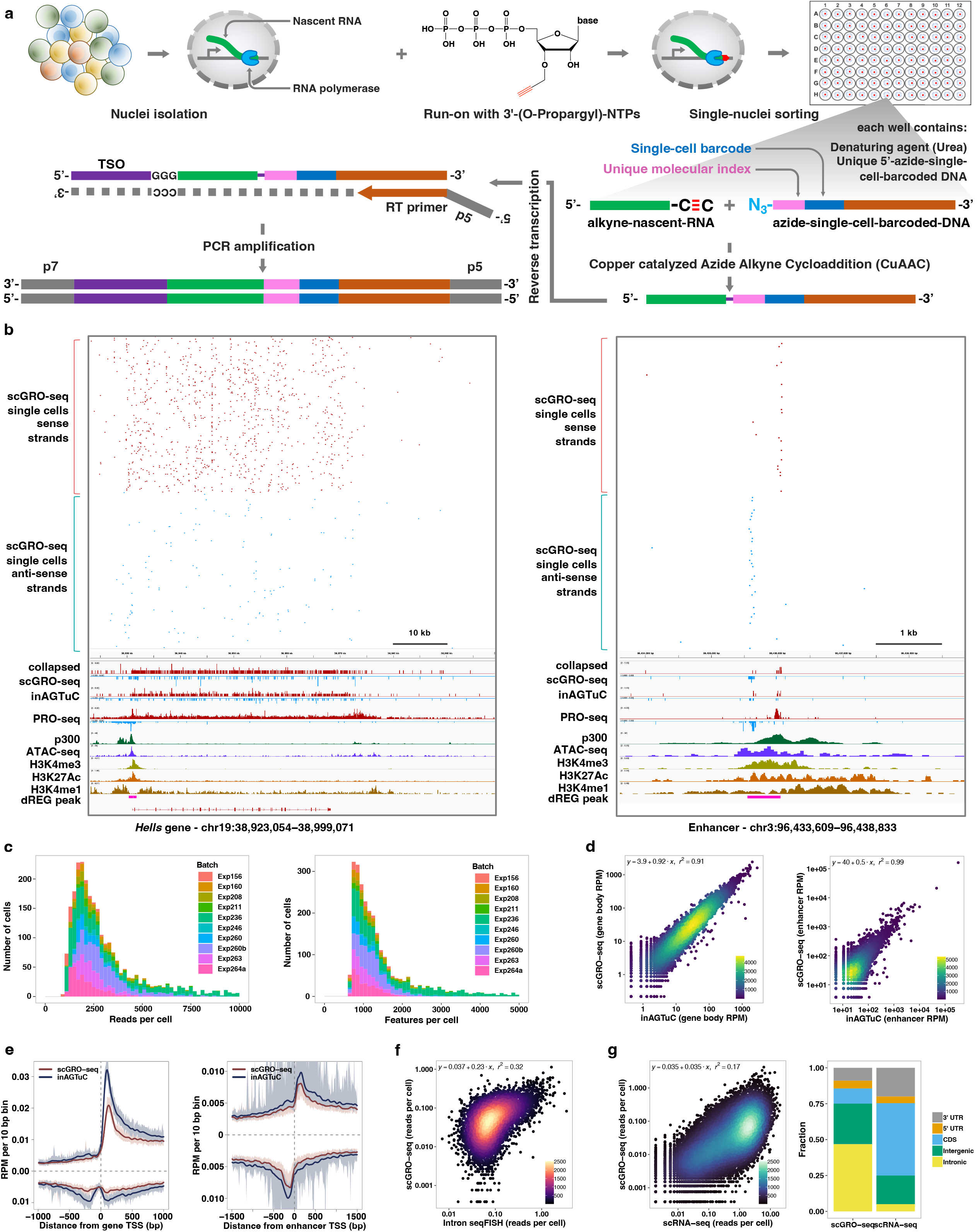
Single-cell nascent RNA sequencing depicts genome-wide nascent transcription. **a,** A summary of scGRO-seq workflow. **b,** Representative genome-browser screenshots showing scGRO-seq reads at a single-cell resolution, the aggregate scGRO-seq level, inAGTuC, PRO-seq, and chromatin marks around a gene (left) and an enhancer (right). **c,** Distribution of scGRO-seq reads per cell (left) and features per cell (right). **d,** Correlation between aggregate scGRO-seq and inAGTuC reads per million sequences in the body of genes (left, n = 19,961) and enhancers (right, n = 12,542). **e,** Metagene profiles of scGRO-seq compared with inAGTuC reads per million per 10 base pair bins around the TSS of genes (left, n = 19,961) and enhancers (right, n = 12,542). The line represents the mean, and the shaded region represents the 95% confidence interval. **f,** Correlation between scGRO-seq and intron seqFISH reads per cell in the body of genes used in intron seqFISH (n = 9,666). **g,** Correlation between scGRO-seq and scRNA-seq reads per cell in the body of genes (left, n = 19,961) and the distribution of reads in various genomic regions (right).

The scGRO-seq recapitulates the inAGTuC and PRO-seq profiles at both genes and enhancers (Fig. 1b) and, for the first time, provides a comprehensive map of nascent transcription in individual cells. We captured an average of 3,665 reads and 1,503 features (genes and enhancers) per cell (Fig. 1c), and pseudo-bulk scGRO-seq counts from collapsed single cells correlate well with bulk counts from inAGTuC (Fig. 1d). The sequencing depth analysis indicated the possibility of more reads and features discovery per cell with further development of technology and deeper sequencing (Extended Data Fig. 7b). However, scGRO-seq is less efficient in capturing nascent RNA from promoter-proximal pause sites. We attribute this to the reduced run-on efficiency of paused Pol II in the absence of a high concentration of strong detergent^27^. This difference in promoter-proximal run-on efficiency is reflected in a lower correlation between scGRO-seq and PRO-seq libraries (Extended Data Fig. 7c), as well as in the metagene profiles around the TSS of genes and enhancers (Fig. 1e & Extended Data Fig. 7d).

After confirming that scGRO-seq recapitulates the bulk nascent RNA sequencing methods, we benchmarked scGRO-seq against other RNA-based single-cell assays. The closest single-cell method that probes nascent transcription is intron seqFISH - a multiplexed single-molecule *in situ* nascent RNA hybridization and imaging method^28^. We confirm that the correlation between scGRO-seq and intron seqFISH is similar to the correlation reported between intron seqFISH and GRO-seq (Fig. 1f). In contrast, scGRO-seq poorly correlates with scRNA-seq (Fig. 1g, left), likely due to a combination of increased mRNA stability and different capture methods^29^. Nevertheless, as expected, scGRO-seq reads are more likely to be intronic or intergenic than scRNA-seq reads (Fig. 1g, right). Overall, the suite of genomic assays presented here utilizes a novel biochemical approach to provide a snapshot of genome-wide transcription at various cell resolutions, including individual cells.

### Direct measurement of transcription burst kinetics

Transcriptional kinetic estimates primarily come from low-throughput live-cell imaging or fluorescent *in situ* hybridization in fixed cells^30–32^. The recently developed Intron seqFISH method is limited to predefined gene targets, requires specialized probes, and assumes that all intronic RNAs have the same kinetic fate. Next-generation sequencing (NGS) based approaches are comprehensive and technically more accessible. However, the current methods measure polyadenylated mRNA from single cells^33^ and fit a simple two-state mathematical model to infer transcriptional kinetics^18^. Bridging this gap, scGRO-seq combines high-throughput measurement of transcription with NGS, thereby enabling the detection of transcribing RNA polymerases genome-wide at single nucleotide resolution (Fig. 2a).

**Fig. 2.**
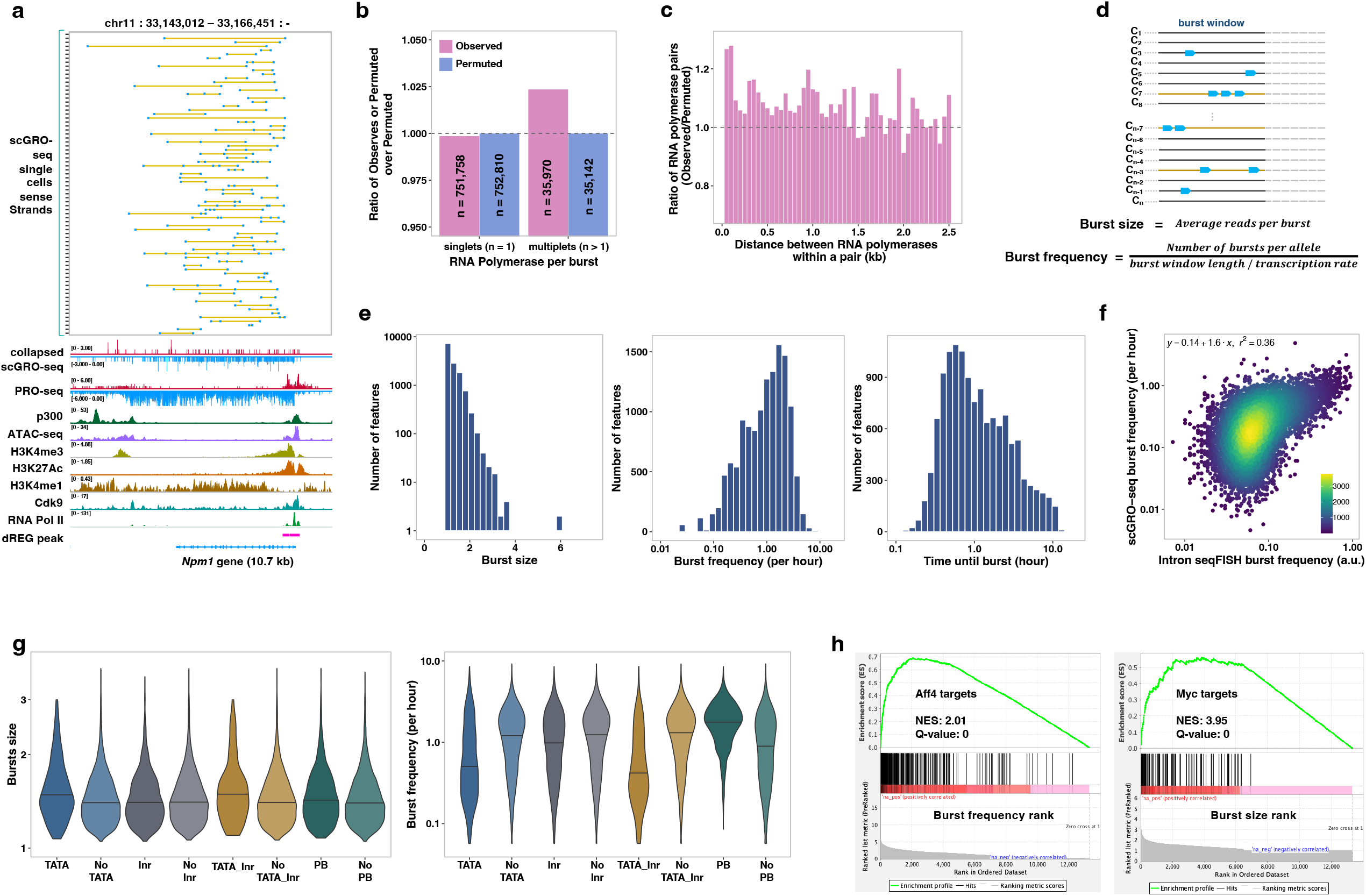
Inference of transcription kinetics using scGRO-seq. **a,** Single-cell view of multiplet RNA Polymerase (blue dots) in *Npm1* gene. A yellow line connects RNA polymerases within the same cell. Randomly sampled 75 single cells containing more than one RNA Polymerase are shown on top, followed by the aggregate scGRO-seq, PRO-seq, chromatin marks, and transcription-associated factors profiles. **b,** Ratio of observed or permuted burst sizes compared against the average burst sizes from 200 permutations. **c,** Ratio of the observed distance between consecutive RNA polymerases in the first 10 kb of gene-bodies in individual cells against the permuted data. Distances up to 2.5 kb are shown in 50 bp bins. **d,** Illustration of the model for direct inference of burst kinetics from scGRO-seq data. **e,** Histogram of burst size (left), burst frequency (middle), and duration until the next burst (1/burst frequency) (right) for genes that are at least 10 kb long (n = 13,142). **f,** Correlation of burst frequency of genes between scGRO-seq and intron seqFISH. **g,** Effect of promoter elements in burst size greater than 1 (left) and burst frequency (right). Inr is Initiator, and PB is pause button sequences. **h,** Role of transcription factors in determining burst frequency and burst size.

With this new approach, we examined the evidence of bursting *de novo* without prior assumptions by quantifying the incidence of transcribing RNA polymerases. If transcription occurs in bursts, we anticipate a higher occurrence of more than one RNA polymerase per burst (multiplets) than would be expected by chance. By permuting reads among cells while keeping their position unchanged (see Methods), we observed a reduction of singlets (n = 1,052, fdr = 0.05) and a concurrent increase in multiplets (n = 828, fdr = 0) in our data compared to the permuted data, providing evidence for the bursting nature of transcription (Fig. 2b). Based on the approximately 10% capture efficiency of scGRO-seq estimated from comparison with intron seqFISH and measurement of transcriptionally engaged RNA polymerases from mammalian cells^34,35^ (see Methods), the probability of detecting two consecutive RNA polymerases on a gene is 1%. The statistically significant increase in 2.4% multiplets exceeds the expectation by chance. Transcriptional bursting would also result in more closely spaced RNA polymerases than what would be observed by random chance^15^. When examining the distance between multiplets, we indeed observed enrichment of closely spaced RNA polymerases (p < 0.05, two-sample KS test) (Fig. 2c & Extended Data Fig. 8a), further strengthening the evidence of bursting.

With confidence in scGRO-seq’s ability to discern bursting, we directly measured the burst kinetics using scGRO-seq counts and their genomic positions. We estimate burst size as the average number of RNA polymerases per burst and burst frequency as the number of bursts per allele per unit of time required for RNA polymerase to traverse through the burst window (Fig. 2d), corrected for capture efficiency (see Methods). We considered genes longer than 11 kb (n = 13,564) and excluded 500 bp regions at either end that are known to harbor paused polymerases^36^, using the remaining 10 kb as the burst window. We assigned reads to a single allele based on the evidence of alleles in mESCs bursting independently generating monoallelic RNA^37,38^. With an average RNA Polymerase II elongation rate of 2.5 kb/min^39^, using a 10 kb region limits the burst detection window to four minutes. This short burst window is consistent with bursts from one allele and aligns with previous reports^12,40^. We simulated kinetic measurements on synthetic data to validate the model’s accuracy and observed robust performance (Extended Data Fig. 8b). We estimated the kinetic parameters of transcriptional bursts for expressed genes (Fig. 2e, Table 1). The mean duration of approximately 2 hours until the next burst in scGRO-seq matches the 2 hours of the global nascent transcription oscillation cycle reported in intron seqFISH. Burst frequency from scGRO-seq correlated well with intron seqFISH (Fig. 2f), and the correlation is more robust for genes with a higher burst frequency. However, we observed a poor correlation between burst frequencies from scGRO-seq and scRNA-seq data, as well as between intron seqFISH and scRNA-seq data (Extended Data Fig. 8c), highlighting potential limitations in the kinetic estimates derived from mature transcripts. Using the burst parameters estimated from scGRO-seq, we again tested our model by simulation and observed robust performance (Extended Data Fig. 8d). Interestingly, no relationship was observed between burst size and burst frequency (Extended Data Fig. 8e), and the gene length did not impact kinetic estimates (Extended Data Fig. 8f). Also, the burst frequencies calculated from 10 kb and 5 kb burst windows showed strong agreement (Extended Data Fig. 8g), confirming the reliability of burst kinetics calculation from scGRO-seq.

**Table 1.** Burst sizes and frequencies of transcribed genes using 5 kb or 10 kb burst windows.

Core promoter elements are known to modulate burst parameters ^18,41,42^. We observed a significant variation in core promoter elements with burst kinetics (Fig. 2g). Specifically, genes with the TATA element exhibited a larger burst size than genes lacking it, and the presence of the Initiator sequence further increased the burst size. The higher burst size but lower burst frequency of genes with TATA elements agree with previous findings^43^. In contrast, highly paused genes, which contain an enriched sequence called pause button at the promoter-proximal paused sites^44^, exhibit higher burst frequency than the remaining genes. The observed difference in burst kinetics among genes with TATA element or promoter-proximal pausing corroborates a previous report of tissue-specific genes exploiting the difference in bursting properties for differential expression across tissues^41^.

Transcription factors are also believed to regulate burst kinetics. Using a curated transcription factor binding database^45,46^, we examined the effect of transcription factors in burst parameters. Gene set enrichment analysis indicated that some transcription factors regulate burst size, and others regulate burst frequency (Table 2). Myc and Aff4 are examples of each category. We found that the genes bound by Myc have larger burst sizes. In contrast, Aff4 target genes are enriched for higher burst frequencies (Fig. 2h). Our observation supports a previous report where Myc increased the burst size by increasing the burst duration^47^, and the association of the Aff4 transcription factor correlated with burst frequency^48^. Overall, we show that the direct and comprehensive observation of transcription using scGRO-seq facilitates the study of transcription kinetics at the single-cell level.

**Table 2.** Gene Set Enrichment Analysis of burst size and burst frequency.

### Cell-cycle inference from non-polyadenylated replication-dependent histone genes

Investigating gene programs during cell-cycle stages is essential for understanding biology and diseases^49^. While scRNA-seq relies on mature transcripts of marker genes to determine the cell-cycle state, the time required for mRNA processing, export, and accumulation introduces a time lag. It also fails to detect the replication-dependent histone genes - the best characterized cell-cycle phase-specific genes exclusively transcribed during the S phase^50^ -due to the lack of polyadenylation^51^. scGRO-seq detects active transcription of replication-dependent histone genes in the histone locus body (Extended Data Fig. 9a) that can be used to classify cells in the S phase. For G1/S and G2/M phase-specific genes, we used a set of transcriptionally characterized genes from an RNA velocity and deep learning study of mESCs^52^. Hierarchical clustering based on the expression of these three cell-cycle phase-specific genes revealed three significant clusters of individual cells (Fig. 3a).

**Fig. 3.**
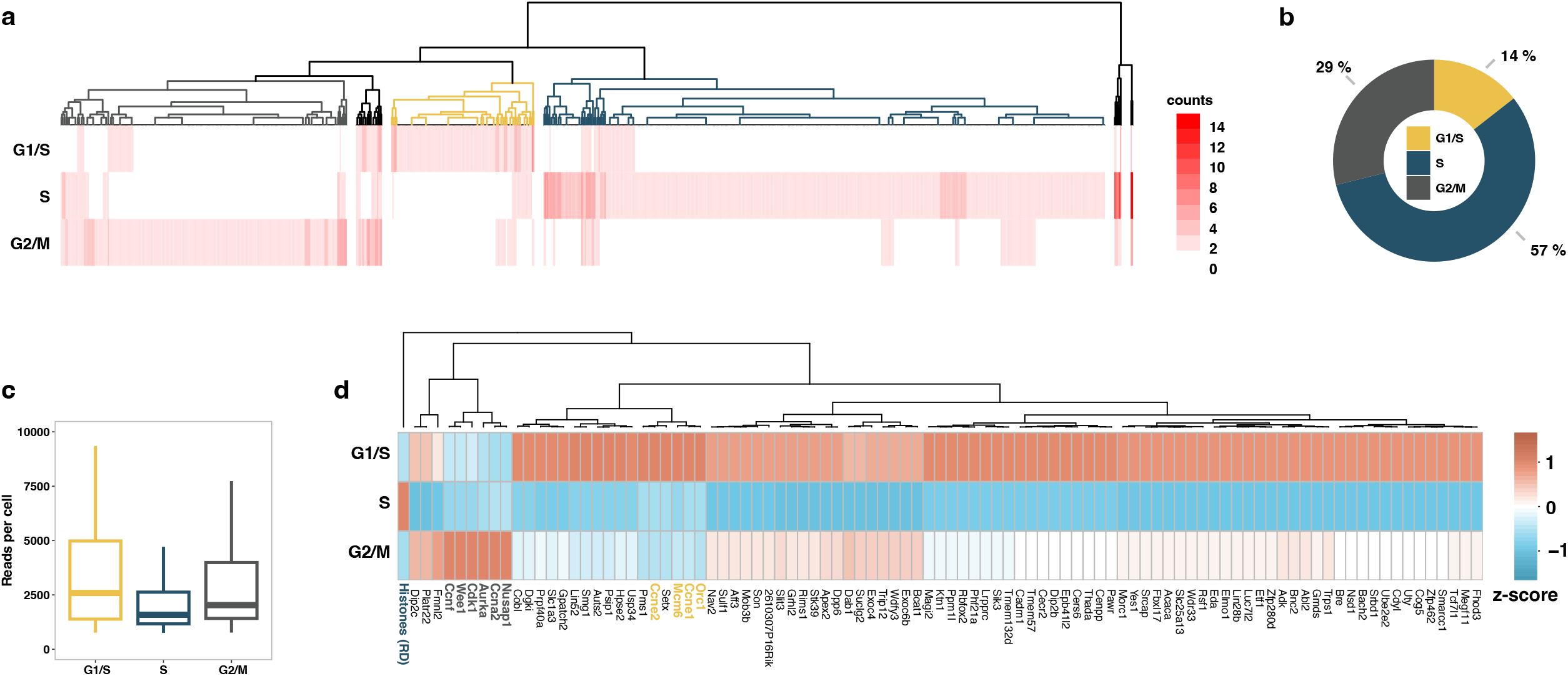
Cell-cycle inference by non-polyadenylated replication-dependent histone genes expression. **a,** Heatmap of hierarchical clustering of single cells representing transcription of G1/S, S, and G2/M specific genes. The dendrogram colors represent cell clusters with cell-cycle phase-specific gene transcription. **b,** Fraction of cells in the three primary clusters distinguished by transcription of G1/S, S, and G2/M specific genes. **c,** Distribution of scGRO-seq reads per cell in the three clusters of cells defined by cell-cycle phase-specific gene transcription. **d,** Differentially expressed genes among the three clusters of cells defined by transcription of G1/S, S, and G2/M specific genes. The genes used to classify cells are denoted in bold and colored. Histones (RD) represent aggregate reads from replication-dependent histone genes.

Mouse embryonic stem cells are known to have a short G1 phase and an extended S phase^53^. De novo classification of mESCs based on the nascent transcription of these newly integrated marker genes recapitulates the cell-cycle phase lengths (Fig. 3b). Notably, cells in the G1/S and G2/M phases exhibit higher transcription levels compared to cells in the S phase (Wilcoxon rank sum test p = 6.3e^-07^ and p = 1.2e^-06^, respectively) (Fig. 3c). We observed an approximately 40% decrease in total transcription when cells transition from G1/S to S phase, with a subsequent 20% increase upon exiting the S phase to G2/M. This observation indicates that transcription continues during DNA replication, albeit at a reduced level. The transition from the G2/M phase to the G1 phase is marked by an increase in transcription^54^, which restores the transcription level observed during the G1 phase, completing the cycle. Analysis of differentially expressed genes in cell-cycle phases also revealed that certain genes restore transcription levels to those observed in the G1/S phase as they transition from the S to the G2/M phase, while others regain partial transcription (Fig. 3d, Table 3). At the same time, some do not recover their transcription until exiting from G2/M to G1/S. By quantifying the active transcription of non-polyadenylated histone genes and a small subset of marker genes, scGRO-seq reveals a dynamic transcription program throughout the cell-cycle.

**Table 3.** Differentially expressed genes among G1/S, S, and G2/M phases of the cell-cycle were identified by the “FindAllFeatures” function of Seurat^1^ (single-cell analysis package).

### Co-transcription of functionally coupled genes

Co-expression of functionally related genes, as measured by accumulated mRNA, is widely reported^55^. However, assessing whether these genes are transcriptionally synchronized in steady-state has been challenging. By utilizing nascent transcription within the first 10 kb of the gene body, thereby limiting the co-transcription detection window to four minutes, we calculated the pairwise Pearson correlation between expressed genes (Fig. 4a). Gene pairs with a correlation coefficient greater than 0.1 and a q-value of less than 0.05, and an empirical false-discovery rate of less than 5% from 1000 permutation were considered co-transcribed (Table 4). These stringent criteria controlled for sampling biases and other confounding effects. We identified only 0.1% of the 112,807,710 candidate gene pairs (n = 137,418) as co-transcribed. We generated a graphical network from these statistically significant pairs, identifying 59 modules (genes per module > 10) of co-transcribed genes. This gene-gene transcriptional correlation could reflect common temporal gene activation by a transcription factor or mechanistic coupling of transcription activation by clusters of genes separated across regions of chromosomes.

**Fig. 4.**
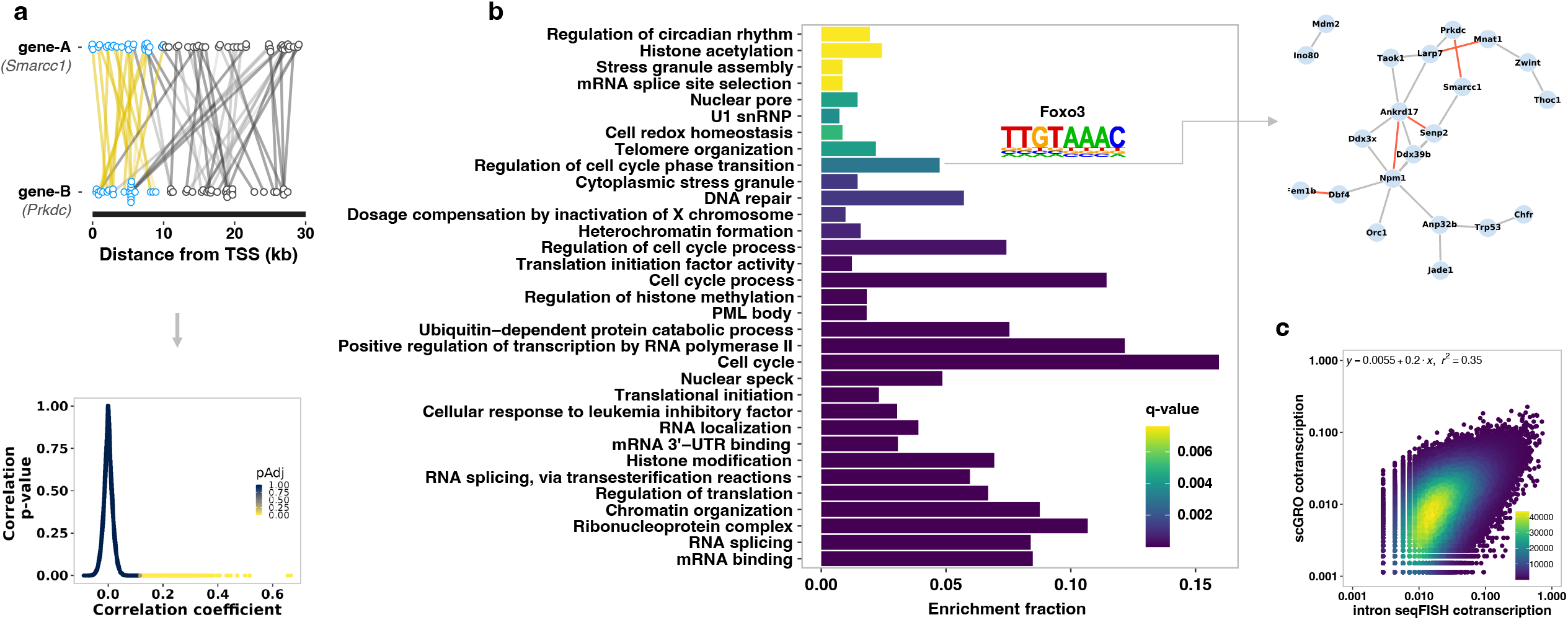
Coordinated transcription of functionally related genes. **a,** A pair of co-transcribed genes (top). Reads within the first 10 kb of the gene pair (blue circle) expressed in the same cells are connected by a yellow line. Reads beyond the first 10 kb (gray circles and lines) are not used in gene-gene correlation. Pair-wise Pearson correlation was calculated from a binarized genes by cells matrix. The relationship between the Pearson correlation coefficient, uncorrected p-value, and the false discovery rate corrected p-value for pairwise gene-gene correlation is shown (bottom). **b,** Gene ontology terms enriched in co-transcribed gene modules. The transcription factor motif enriched in the promoters of the genes associated with the GO term and the co-transcribed genes that contributed to the enrichment of the GO term is shown as an example on the right (red line indicating rho > 0.15). A complete list of GO terms and the co-transcribed genes contributing to the enrichment of the GO terms is provided in Table 5. **c,** Correlation of co-transcription of significantly co-transcribed gene pairs (n = 164,380) between scGRO-seq and intron seqFISH. Axes represent the fraction of cells in which a gene pair is co-transcribed.

**Table 4.** Correlated gene-gene pairs.

**Table 5.** Non-redundant Gene Ontology terms and the co-transcribed genes that contribute to the GO terms’ enrichment.

Conducting gene ontology analysis on these co-transcribed modules compared to all transcribed genes, we found enrichment of several related molecular functions, including cell-cycle regulation, RNA splicing, translational control, DNA repair, and circadian rhythm (Fig. 4b, Extended Data Fig. 9b, and Table 5). By scanning the promoters of co-transcribed genes, we discovered an enrichment of known transcription factor motifs, such as Foxo3 enriched in the promoters of co-transcribed genes associated with the “Regulation of cell-cycle phase transition” gene ontology term. A previous study has shown that Foxo3, in coordination with the DNA replication factor Cdt1, is critical in regulating cell cycle progression^56^. We compared the co-transcription patterns of gene pairs obtained from scGRO-seq with those from intron seqFISH, and the results revealed concordant co-transcription (Fig. 4c). This high throughput and unique capability of scGRO-seq to directly examine transcriptional coordination between any gene pair or network of genes provides valuable insights into the functional organization of the genome.

### Spatiotemporal coordination of enhancer-gene co-transcription

Regulation of gene expression by distal regulatory elements is an area of broad interest. scGRO-seq captures transcripts from both genes and active enhancers, allowing the measurement of co-activation in single cells. We analyzed scGRO-seq reads within the first 10 kb of genes and at least 3 kb on each strand transcribing outwards around enhancers (see Methods). We excluded 500 bp regions around the TSS of genes and enhancers to exclude paused polymerase. We also included clusters of enhancers known as super-enhancers (SEs) that do not overlap with gene regions^20^.

We identified enhancer-gene pairs that exhibit significant pairwise Pearson correlations on the same chromosomes. Out of 2,718,382 test pairs, 1.8% (n = 49,143) passed the threshold criteria of correlation coefficient, q-value, and empirical false-discovery rate from 1000 permutations (Table 6). We observed a significant enrichment (two-sample KS test, p-value = 5e^-07^) of enhancer-gene co-transcription within 100 kb of each other (Fig. 5a), with SE-gene pairs showing even stronger enrichment (two-sample KS test, p-value = 1.7e^-09^) compared to uncorrelated pairs (Extended Data Fig. 10a). When examining functionally related genes clustered together on the same chromosome^57^, we found multiple enhancers correlated with each gene (Extended Data Fig. 10b), suggesting the possibility of an increased concentration of transcription machinery locally through condensate formation, or a further manifestation of cell-cycle regulation.

**Fig. 5.**
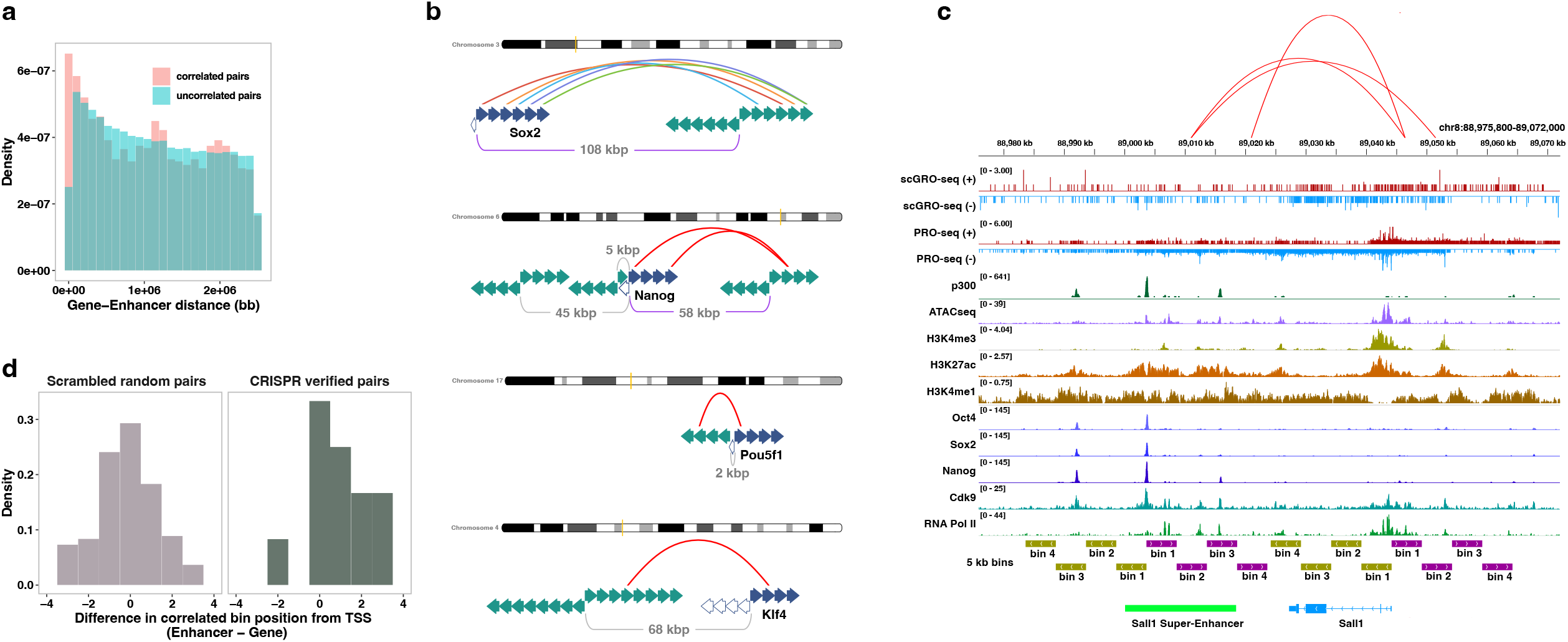
Spatial and temporal coordination between genes and enhancers. **a,** Distance between correlated and non-correlated enhancer-gene pairs within 2.5 Mb of each other. **b,** Co-transcription between pluripotency genes (filled blue arrows indicate sense gene bins, empty blue arrows indicate antisense gene bins) and their enhancers (represented by green arrows, and the arrow directions indicate sense and antisense directions). Correlated full-length enhancer-gene pairs (Sox2 and Nanog) are shown with purple distance bars. For finer time resolution correlation, features are extended up to the end of the transcription signal and divided into 5 kb bins. Correlated bins are represented by a red arch, except for Sox2 and its distal enhancer bins, which are shown in different colors for visual aid. **c,** Co-transcription between the Sall1 gene and its CRISPR-verified SE. Correlated SE-gene bins are denoted by arches, **d,** Summary of correlated bin positions in CRISPR-verified SE-gene pairs. Scrambled random pairs serve as a control.

**Table 6.** Correlated enhancer-gene pairs.

We investigated a set of validated enhancers known to regulate pluripotency transcription factors^58–61^. We observed statistically significant correlations between the transcription of the Sox2 and Nanog genes and their distal enhancers (Extended Data Fig. 10c). The lack of correlation between the Oct4 and Klf4 enhancer-gene pairs could be due to a time delay in the transcription of genes and enhancers. If enhancers and their target genes are temporally coupled and co-transcribe, we speculated that the co-transcription of the pair could be even more prominent at finer temporal resolution. To test this, we divided enhancers and genes into 5 kb bins (representing a 2-minute transcription window) and found that at least one enhancer bin correlated significantly with its target gene for all four genes (Fig. 5b). Intriguingly, the correlated enhancer bin generally appeared further from its TSS than the gene bin, implying that enhancer transcription may initiate prior to promoter transcription.

To test the enhancer-gene timing hypothesis, we examined a set of seven non-intronic mESC super-enhancers validated by CRISPR perturbation^62^. The CRISPR knock-out of Sall1 SE reduced Sall1 expression by 92%, and we found a correlation between multiple enhancer bins and this gene (Fig. 5c). Overall, 4 out of 7 SE-gene pairs showed correlations of at least one bin. Interestingly, we observed that in most cases, enhancer transcription began earlier or around the same time as the transcription of their target genes (Fig. 5d). This temporal pattern could have mechanistic implications for enhancer-gene regulation. However, any conclusions will require a much deeper data set. Nevertheless, our findings offer a glimpse into the possibility of temporal order in enhancer-gene transcription.

## Discussion

We developed scGRO-seq to enable the assessment of co-transcription and prediction of enhancer-gene regulatory networks in their native context. By reporting the activity of genes and distal regulatory elements - and thus the functional consequences of transcriptional signals and networks - scGRO-seq is inherently multimodal for understanding transcription regulation at an unprecedented level. We illustrate these advantages by determining burst size and frequency for expressed genes, transcription levels during cell cycle phases, and genome-wide gene-gene and enhancer-gene co-transcription detection. We restricted this study to mESCs for comparison with large available data sets for validation.

The current scGRO-seq methodology has its limitations. The preservation of nuclear integrity, achieved through a low sarkosyl concentration, fails to promote the run-on of RNA polymerases in the pause complex, thereby limiting the detection of promoter-proximal paused polymerases. The read depth and cell numbers limit our analyses of burst kinetics and synchrony of gene-gene and enhancer-gene pairs. Improved efficiency in future iterations will facilitate a more precise evaluation of these phenomena.

scGRO-seq is also limited by the abundance of nascent RNA per cell at any given time, which is considerably lower than that of mature mRNA. Nascent RNA detection requires technology not dependent on a polyadenylated terminus, initially raising doubts about the feasibility of nascent RNA sequencing in single cells^63^. However, implementing highly efficient CuAAC has overcome this limitation, enabling the capture of approximately 10% of nascent RNA with current protocols. To streamline the process and ensure compatibility with future automation, we optimized the biochemical steps by replacing multiple rounds of nascent RNA purification and nucleic acid ligation with click-chemistry. Further adaptations, including high-throughput droplet encapsulation and enhanced capture efficiency, will extend the applicability of our scGRO-seq method in both research and clinical settings.

For clinical specimens, especially for challenging tissues like the brain and pancreas that contain high levels of RNase, isolation of nuclei is more feasible than obtaining intact single cells. Single-cell methods like sNuc-Seq^64^ profile polyadenylated RNA inside the nucleus of such tissues, painting an incomplete view of single-cell gene expression. In contrast, the entire scGRO-seq substrate is present inside the nucleus. Furthermore, the compatibility of CuAAC-based nascent RNA sequencing methods with bulk, small input samples, and single cells makes them desirable methods for clinical investigations. The adaptability and efficiency of scGRO-seq introduce new avenues for investigating transcriptional dynamics and regulatory mechanisms across diverse biological contexts, enriching our understanding of gene expression regulation and its ramifications in physiological and pathological conditions.

## Methods

### scGRO-seq conceptualization

Capturing nascent RNA with sufficient efficiency from single cells for meaningful analysis was deemed challenging^2^. However, recognizing the potential insights into transcription mechanisms that single-cell nascent RNA sequencing could offer, we set out to develop a single-cell version of the GRO-seq method a decade after its use in cell populations^3^. Our efforts were met with two major challenges: selectively capturing a small fraction of nascent RNA among various RNA species within a cell and accurately distinguishing nascent RNAs from individual cells.

The primary limitation we encountered was capture efficiency. The quantity of nascent RNA from transcribing RNA polymerases in an individual cell, mainly due to transcription’s intermittent nature with short bursts and long latency periods, is significantly lower than the mRNA copies that accumulate over time. Traditional nascent RNA capture methods yield only a meager number of nascent RNAs from single cells. Miniaturizing GRO-seq using strategies derived from scRNA-seq was not feasible since nascent RNA lacks the consensus polyadenylation sequence used in RNA-seq. Instead, GRO-seq and related methods selectively label nascent RNA in bulk cells using modified nucleotides and employ single-stranded RNA-RNA ligation with PCR handles on both ends. This ligation process proved unsuitable for scGRO-seq due to its low efficiency and the need for nascent RNA purification before ligation, which risks depleting the already scarce nascent RNA from single cells.

To overcome these challenges, we devised a strategy involving a) labeling nascent RNA in cells and b) attaching single-cell barcodes to the labeled nascent RNA without requiring purification from other cellular RNA. After exploring several approaches without success, we turned to click-chemistry, specifically CuAAC (copper(I)-catalyzed azide-alkyne cycloaddition). We speculated that by sourcing or synthesizing CuAAC-compatible chain-terminating nucleotide triphosphate analogs and performing nuclear run-on with the modified nucleotides to label nascent RNA selectively, we could label nascent RNA from individual cells with 5’-azide-single-cell-barcoded (5’-AzScBc) DNA with a PCR handle. Then, we could pool the barcoded nascent RNA from multiple cells for selective reverse transcription in the presence of a template switching oligo (TSO) and subsequent PCR amplification for sequencing.

To successfully implement this strategy, we identified three critical biochemical hurdles to address. First, we needed to demonstrate the ability of native RNA polymerase to incorporate 3’O-Propargyl-NTPs during nuclear run-on reactions. Second, preserving the intactness of nuclei during the run-on reaction was essential to enable the separation of individual nuclei for single-cell barcoding. Finally, we had to confirm the ability of reverse transcriptase to traverse the triazole ring junction formed during CuAAC. Successful resolution of the first and third hurdles would pave the way for CuAAC-based nascent RNA sequencing in cell populations while overcoming the second hurdle would establish the foundation for single-cell GRO-seq.

### Development of AGTuC

To develop a nascent RNA tagging method suitable for capturing a small fraction of RNA from single cells, we initiated our approach by focusing on a cell-population-based strategy. Our aim was to develop an enhanced nascent RNA tagging method that optimally integrates selective labeling and single-cell barcode tagging, bypassing the need for RNA purification. Among the tested methods, we identified click-chemistry as the most suitable option due to its high selectivity, efficiency, robustness in diverse experimental conditions, cost-effectiveness, and speed. Our goal was to selectively label nascent RNA through a nuclear run-on reaction, conjugate a single-stranded DNA PCR handle (that can accommodate a single-cell barcode for future use in single-cell analysis), reverse transcribe the RNA-DNA conjugate, and prepare a next-generation sequencing (NGS) library.

To achieve single-nucleotide resolution of transcribing polymerases and efficient reverse transcription, we identified two click-chemistry compatible, chain-terminating nucleotides with relatively small functional group - 3’-(O-Propargyl)-ATP and 3’-Azido-3’-dATP (Extended Data Fig. 1a). Nascent RNA labeled with 3’-(O-Propargyl)-NTPs forms a 1,4- disubstituted 1,2,3-triazole junction with azide-labeled DNA through copper-catalyzed azide-alkyne cycloaddition (CuAAC), whereas nascent RNA labeled with 3’-Azido-3’- dNTPs forms a slightly bulkier junction with dibenzocyclooctyne (DBCO) labeled DNA via Strain-Promoted Alkyne-Azide Cycloadditions (SPAAC) (Extended Data Fig. 1b). Nuclear run-on with 3’-(O-Propargyl)-ATP and CuAAC showed superior efficiency compared to 3’- Azido-3’-dATP and SPAAC (Extended Data Fig. 1c).

To convert the clicked RNA-DNA conjugate to cDNA, we tested eight different reverse transcriptase enzymes, varied the temperature and duration of reverse transcription, and evaluated three template-switching oligos (Extended Data Fig. 1d-f, some results not shown). Our optimized method, which we named the Assay for Genome-wide Transcriptome using Click-chemistry (AGTuC), was then performed in 5 million mESCs nuclei. AGTuC nascent RNA profiles closely resembled PRO-seq profiles (Extended Data Fig. 2a) and exhibited strong correlations at both gene and enhancer levels (Extended Data Fig. 2b-c). Notably, the AGTuC library protocol involved significantly fewer steps than PRO-seq and could be completed in a single day (Extended Data Fig. 2d). AGTuC is a simpler, faster, and cheaper alternative to GRO-seq and PRO-seq for nascent RNA sequencing from cell populations.

### Development of inAGTuC

To adapt CuAAC-mediated nascent RNA sequencing to single cells, we explored the feasibility of performing AGTuC in single cells. Implementing AGTuC at the single-cell level presented challenges, as the nuclear run-on reaction with 0.5% sarkosyl disrupts the nuclear membrane before cell barcodes could be attached during the post-run-on CuAAC step, leading to unintended mixing of nascent RNA from different cells. One potential solution was to perform AGTuC in single tubes, which would prevent nascent RNA mixing. However, this approach requires RNA purification after the run-on reaction, but purification results in further depletion of exceedingly low amounts of nascent RNA in single cells. Alternatively, omitting RNA purification would lead to an abundance of 3’-(O-Propargyl)-NTPs supplied in huge excess during the run-on reaction, which could outcompete 5’-AzScBc DNA during CuAAC.

To address this challenge, we developed intact-nuclei AGTuC (inAGTuC) - a novel strategy that enables labeling nascent RNA with 3’-(O-Propargyl)-NTPs while preserving nuclear integrity. This approach overcomes the issues associated with nascent RNA mixing prior to single-cell barcoding. We hypothesized that performing the run-on reaction without disrupting the nuclear membrane would facilitate the easy removal of excess nucleotides through a few centrifugation and resuspension steps while retaining propargyl-labeled nascent RNA within the nuclei. This approach would yield clean nuclei with labeled nascent RNA, free from excess reactive nucleotides, which could be compartmentalized with 5’-AzScBc DNA for CuAAC. We could minimize further RNA loss by pooling and processing the single-cell-barcoded nascent RNA from multiple cells.

To achieve an efficient run-on reaction, PRO-seq and AGTuC disrupt the polymerase complex with 0.5% sarkosyl detergent, of which nuclear membrane lysis is collateral damage. We sought to identify the lowest sarkosyl concentration that maintains nuclear membrane integrity while maximizing run-on efficiency and found that a 20x reduction in sarkosyl concentration preserved nuclear intactness, with only a 20% reduction in run-on efficiency (Extended Data Fig. 3a-b). To maximize the nascent RNA capture efficiency, we optimized the molecular crowding effect of PEG 8000 and the ratio of Cu(I) to CuAAC accelerating ligand BTTAA (Extended Data Fig. 3c). Although a low sarkosyl concentration preserves nuclear integrity, it also retains the RNA polymerase complex intact, thereby shielding the propargyl-labeled 3’ end of nascent RNA from reacting with 5’-AzScBc DNA. We investigated nascent RNA release from the RNA polymerase complex using common denaturants and found 6 M urea and Trizol to be efficient (Extended Data Fig. 3d). However, the denaturant in Trizol hindered CuAAC reaction (Extended Data Fig. 3e). Remarkably, urea also offered the added benefit of retaining the RNA-DNA conjugate in the aqueous phase during Trizol clean-up to remove PEG 8000 from the CuAAC reaction (Extended Data Fig. 3f). For reaction clean-up, we assessed various methods, finding cellulose membrane to be effective in removing CuAAC reagents (Extended Data Fig. 3g), while silica matrix columns performed well in retaining RNA and ssDNA (Extended Data Fig. 3h). Subsequently, we evaluated DNA polymerase for library preparation and DNA size-selection methods (Extended Data Fig. 3i-j).

Considering the goal of working with single cells, we performed inAGTuC with cell numbers between 5 million used in AGTuC and one cell planned for scGRO-seq. Specifically, we placed 100 to 1000 intact nuclei in each well of a 96-well plate containing Urea. Nascent RNA in each well was barcoded with a unique 5’-AzScBc DNA by CuAAC and pooled from the 96 wells, and a sequencing library was prepared as in AGTuC. The inAGTuC libraries exhibited similar profiles in gene bodies compared to PRO-seq and AGTuC. However, they could not capture the paused peaks at the 5’ end of genes and enhancers (Extended Data Fig. 4a-c). This observation is consistent with the need for a higher sarkosyl concentration for efficient run-on of paused Polymerase complexes^4^. The four inAGTuC libraries correlated well with each other (Extended Data Fig. 4d), with the potential to discover more insights with deeper sequencing (Extended Data Fig. 4e-f). Despite only partially capturing nascent RNA from a paused complex, the inAGTuC libraries correlated well with AGTuC and PRO-seq (Extended Data Fig. 4g).

To systematically characterize the compatibility of inAGTuC with even fewer cells, we prepared four inAGTuC libraries in a 96-well plate, with 12 cells per well (cpw), 120 cpw, and 1200 cpw. We also included a 1200 cpw plate, omitting Cu(I) as a negative control. Despite lower coverage, the inAGTuC library with 12 cpw (total ∼1,000 cells) successfully captured the overall nascent RNA profile. It exhibited a good correlation with 120 cpw (total ∼10,000 cells) and 1200 cpw (total ∼100,000 cells) (Extended Data Fig. 5a-c).

### 3’-(O-Propargyl)-Nucleotide synthesis

For this study, several CuAAC-compatible nucleotide analogs modified with azide or alkyne functionalities were evaluated. Ultimately, 3’-(O-Propargyl)-NTPs were selected for three main reasons:

a. These analogs lack 3’ hydroxyl groups, making them chain-terminating and enabling single-nucleotide resolution of the 3’-end of nascent RNA.
b. The CuAAC reaction produces a compact junction due to the presence of a single carbon bond between the sugar group of the nucleotide and the propargyl group at the 3’-end position.
c. They are relatively cost-effective compared to biotin-modified nucleotides commonly used in PRO-seq.

3’-(O-Propargyl)-ATP (NU-945) was offered by Jena Biosciences. To complete the set, custom synthesis requests were made for 3’-(O-Propargyl)-CTP (NU-947), 3’-(O-Propargyl)-GTP (NU-946), and 3’-(O-Propargyl)-UTP (NU-948), all of which are now available for purchase from Jena Biosciences.

### Single-cell barcoded DNA adaptors

Over the course of scGRO-seq development, we synthesized three sets of 96 5’-AzScBc DNA from GeneLink. Each design had four components: 5’ azide at the 5’ end, single-cell barcode sequence, unique molecular index (UMI), and PCR handle. The 5’ azide modification was obtained as described before^5^. Briefly, an oligonucleotide with 5’ Iodo-dT was synthesized on solid support by phosphoramidite oligo synthesis, and the Iodo group was replaced with azide by reacting with sodium azide at 60°C for 1 hour.

During scGRO-seq development, three sets of 96 5’-AzScBc DNA were synthesized by GeneLink. Each design encompassed four components: a 5’ azide positioned at the 5’ terminus, a 10-12 nucleotide sequence for the single-cell barcode, a 4-6 nucleotide sequence for unique molecular index (UMI), and a PCR handle. The 5’ azide modification was obtained following a previously described method^5^. Specifically, an oligonucleotide containing 5’ Iodo-dT was synthesized via solid-support phosphoramidite oligo synthesis, and subsequent replacement of the Iodo group with an azide group was achieved through a reaction with sodium azide at 60°C for 1 hour. The sequences of three different 5’-AzScBc DNA are listed below:

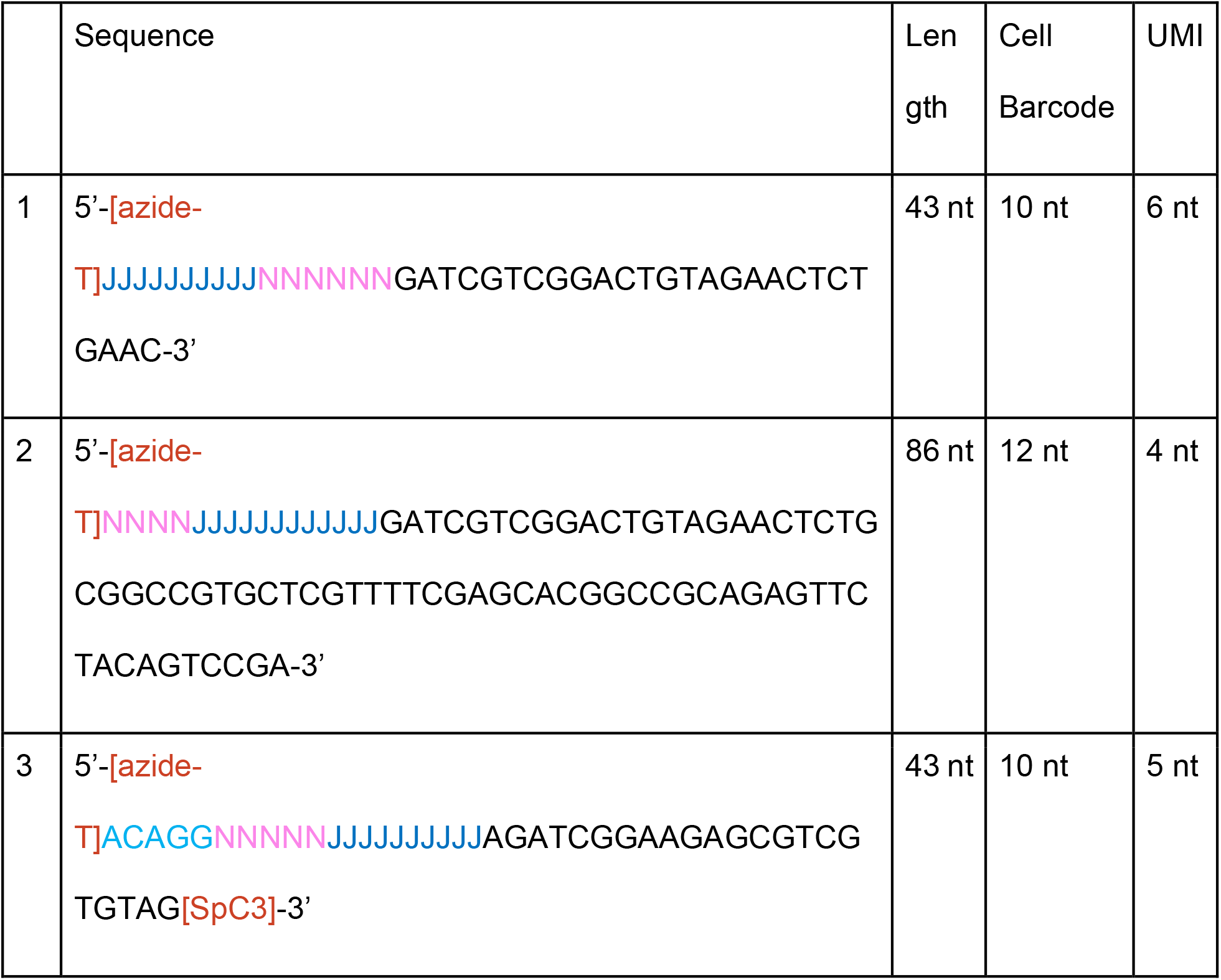

Js represent unique cell barcodes, Ns represent Unique molecular index, and SpC3 denotes a three-carbon spacer.

The hairpin structure of the 86-nucleotide 5’-AzScBc DNA is formed through self-folding. The reverse transcription process is initiated using the 3’ end of the oligo, which serves as a built-in primer.

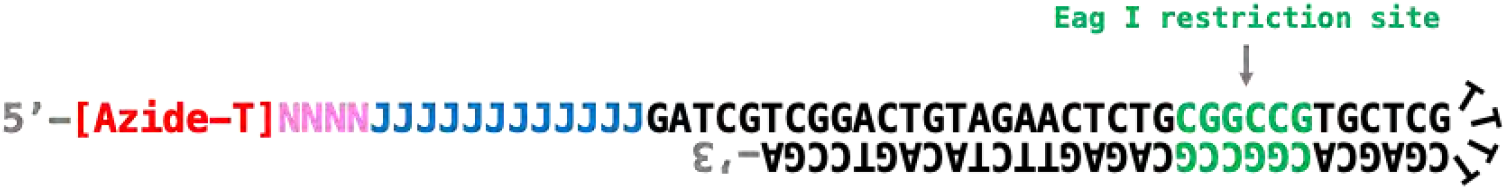

The design ensures a 1:1 stoichiometry between the PCR handle and RT primer, minimizing mispriming and non-specific amplification during reverse transcription. The folded hairpin structure also generates a restriction site for the Eag I enzyme, which is digested before PCR amplification.

Undesired extension by reverse transcriptase is effectively prevented by a three-carbon spacer at the 3’ end of the 43-nucleotide 5’-AzScBc DNA^6^. This spacer harbors a 5-nt ACAGG sequence after the azide-dT at its 5’ end. During RT, the extension of primers annealing to unclicked 5’-AzScBc, the addition of non-templated CCC, and template switching oligo (TSO) incorporation results in undesired cDNA that are preferred substrates for PCR amplification. If unaddressed, these amplicons can overwhelm the sequencing library. The ACAGG sequence plays a critical role in depleting these PCR amplicons.

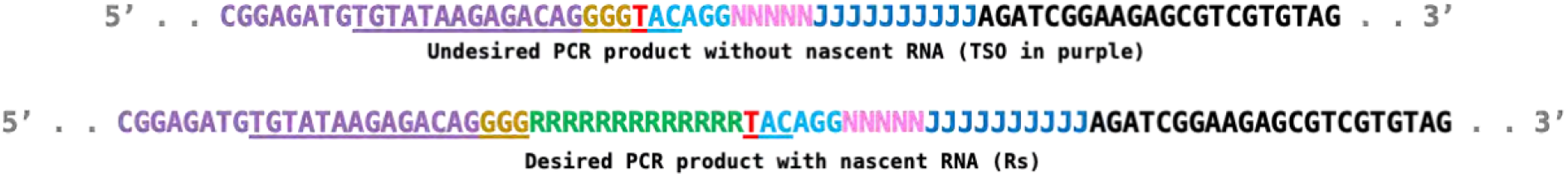

A previously described method named DASH uses recombinant Cas9 protein and gRNA complex to digest and deplete undesired dsDNA^7^. The ACAGG sequence is necessary to generate a guide RNA target sequence in the undesired PCR amplicons (underlined sequence). In PCR amplicons formed between nascent RNA and 5’-AzScBc DNA, the complementation of gRNA is interrupted by the presence of a nascent RNA sequence, making the desired products incompatible with DASH. AGG serves as the protospacer adjacent motif (PAM).

### Cell line

V6.5 mouse embryonic stem cells (mESCs) used in this study were established by Jaenisch laboratory (Whitehead Institute, Massachusetts Institute of Technology) from the inner cell mass (ICM) of a 3.5-day old mouse embryo from a C57BL/6(F) X 129/sv(M) cross.

### Cell Culture

mESCs were cultured in Dulbecco’s Modified Eagle Medium (Gibco, 11995), plus 10% fetal bovine serum (HyClone, SH30070.03), supplemented with 1x penicillin/streptomycin (Gibco, 15140), 1x non-essential amino acids (Gibco, 1140), 1x L-Glutamine (Gibco, 25030), 1x β-mercaptoethanol (Sigma, M6250), and 1000 U/ml leukemia inhibitory factor (Sigma, ESG1107) on tissue culture-treated 10 cm plates (Corning, CLS430167) pre-coated with 0.2% gelatin (Sigma, G1890) prepared in phosphate-buffered saline (Fisher, MT21031CV). Cells were grown at 37°C and 5% CO_2_ and passed with Hepes buffered saline solution (Lonza, CC-5024) and 0.25% Trypsin-EDTA (Gibco, 25200) when 70% confluency was reached (every two days).

### Sample preparation

Tissue culture cells were prepared for nuclear run-on reaction by either nuclei isolation or cell permeabilization as described before^2^. All centrifugation steps were performed at 1000 g for 5 min. Cells were harvested by dumping the tissue culture media, rinsing with PBS, and placing the plates on ice. Cells were scraped while still on ice. The harvest was collected into a 15 ml conical tube and centrifuged at 1000 g for 5 min.

For nuclei isolation, the pellet was resuspended in ice-cold douncing buffer (10 mM Tris-Cl pH 7.4, 300 mM Sucrose, 3 mM CaCl_2_, 2 mM MgCl_2_, 0.1 % Triton X-100, 0.5 mM DTT, 0.1x Halt Protease inhibitor, and 0.02 U/ul RNase inhibitor) and transferred to a 7 ml dounce homogenizer (Wheaton, 357542). After incubation on ice for 5 minutes, the cells were dounced 25 times with a tight pestle, transferred back to the 15 ml conical tube, and centrifuged to pellet the nuclei. The pellet was washed twice in douncing buffer.

For cell permeabilization, the pellet was resuspended in ice-cold permeabilization buffer (10 mM Tris-Cl pH 7.4, 300 mM Sucrose, 10 mM KCl, 5 mM MgCl_2_, 1 mM EGTA, 0.05 % Tween-20, 0.1% NP-40, 0.5 mM DTT, 0.1x Halt Protease inhibitor, and 0.02 U/ul RNase inhibitor). After incubation on ice for 5 minutes, the cells were centrifuged to pellet the nuclei. The pellet was washed twice in the permeabilization buffer.

The washed pellet was resuspended in storage buffer (10 mM Tris-Cl pH 8.0, 5% Glycerol, 5 mM MgCl_2_, 0.1 mM EDTA, 5 mM DTT, 1x Halt Protease inhibitor, 0.2 U/ul RNase inhibitor) at a concentration of 5×10^6^ nuclei per 50 ul of storage buffer, flash-frozen in liquid nitrogen and stored at -80°C. The nuclei and permeabilized cells in the storage buffer can be stored for up to 5 years at -80°C, making them readily available for nuclear run-on experiments.

### Nuclear Run-on with 3’-O-Propargyl Nucleotides

50 ul of 2X nuclear run-on buffer (20 mM Tris-Cl pH 8.0, 10 mM MgCl_2_, 400 mM KCl, 50 uM 3’-(O-Propargyl)-ATP, 50 uM 3’-(O-Propargyl)-CTP, 50 uM 3’-(O-Propargyl)-GTP, and 50 uM 3’-(O-Propargyl)-UTP, 0.05% Sarkosyl, 1 mM DTT, 2X Halt Protease inhibitor, and 0.4 U/ul RNase inhibitor) was prepared per sample and heated to 37°C. Once thawed from -80°C, permeabilized cells or nuclei were added to the heated tube containing nuclear run-on buffer and incubated for 5 min at 37°C with gentle tapping at the incubation midpoint. Permeabilized cells or nuclei were centrifuged at 500 g for 2 min at 4°C, and the supernatant was aspirated off. The pellet was washed three times in 150 ul Resuspension buffer (5 mM Tris-Cl pH 8.0, 2.5 % Glycerol, 2.5 mM MgAc_2_, 0.05 mM EDTA, 1.25 mM MgCl_2_, 60 mM KCl, 3 mM DTT, 0.2X Halt Protease inhibitor, and 0.2 U/ul RNase inhibitor). After the final wash, the permeabilized cells or nuclei were resuspended in a 2 ml Resuspension buffer and passed through a 35 µm nylon mesh (Falcon, 352235).

### Single-cell/nuclei sorting

96-well plates with 2.5 ul 8 M Urea are prepared for single-cell/nuclei sorting using a multi-channel or 96-well pipettor (Avidien MicroPro 300, 30835029). Single cells/nuclei populations characterized by the forward and side scattering are sorted into the 96-well plate containing Urea by fluorescent activated cell sorter (FACS). The sorted plates can be used in CuAAC directly or sealed with aluminum foil or a plastic seal and stored at - 80°C.

### CuAAC

A 96-well plate containing 5’-AzScBc DNA with a unique cell barcode in each well previously synthesized and aliquoted is thawed from -80°C. Sodium Ascorbate, PEG 8000, CuSO_4_, and accelerating ligand BTTAA were prepared and dispensed into each well of the 96-well plate containing 5’-AzScBc DNA. The CuAAC reaction mix was dispensed into individual wells containing single cells in Urea using a multi-channel or 96-well pipette. The final concentration of CuAAC reaction in each well is 30 nM 5’-AzScBc DNA, 800 mM Sodium Ascorbate, 15% PEG 8000, 1 mM CuSO4, and 5 mM BTTAA, and 2.66 M Urea in a 7.5 ul volume. The 96-well plates are sealed, vortexed for 10 seconds in an orbital vortexer, and centrifuged for 1 min at 500 g before incubation for 2 hours at 50°C.

After incubation, the CuAAC reaction was quenched with 5 mM EDTA and pooled from 96 wells into a 1.5 ml Eppendorf tube. PEG 8000 was removed with Trizol. The remaining CuAAC reagents (Sodium Ascorbate, CuSO4, and mM BTTAA) were removed with a centrifugal filter with 3 kDa cellulose membrane (Amicon, 2020-04). The purified RNA was fragmented with 10 mM ZnCl_2_ for 5 min at 65°C.

### Reverse transcription through the triazole link and pre-amplification

Reverse transcription of the clicked RNA-DNA conjugate was performed with highly processive Moloney Murine Leukemia Virus (M-MuLV) Reverse Transcriptase lacking RNase H activity but capable of RNA-dependent and DNA-dependent polymerase activity, non-templated addition and template switching (Thermo Fisher, EP0751). RT reaction (1x RT buffer, 0.5 mM dNTPs, 0.8 U/ul RNase inhibitor, 16% PEG 8000, 1 uM RT primer (except for hairpin forming 5’-AzScBc DNA), and 1 um TSO) was incubated with RNA-DNA conjugate for 2 hours at 50°C. The cDNA was size-selected in 10% denaturing polyacrylamide gel electrophoresis (PAGE) away from the unclicked 5’- AzScBc DNA and empty cDNA formed between the 5’-AzScBc DNA and TSO.

The purified cDNA was PCR amplified for 6 cycles to generate dsDNA with NEBNext Ultra II Q5 High-Fidelity 2X Master Mix (NEB, M0544) and 0.5 uM PCR primers with unique dual index using the following PCR cycles:

|  | Steps | Denature<br>(98°C) | Anneal<br>(60°C) | Extension<br>(72°C) | Storage<br>(4°C) |
| --- | --- | --- | --- | --- | --- |
|  | Initial denaturation | 30 sec | - | - | - |
| 6 cycles | Denaturation | 10 sec | - | - | - |
|  | Anneal |  | 20 sec | - | - |
|  | Extension |  |  | 30 sec |  |
|  | Final extension | - | - | 2 min | - |
| | Storage | - | - | - | ∞ |

### Removal of empty adaptors using DASH

The dsDNA from the pre-amplification of cDNA was subjected to DASH to remove the undesired amplicons formed by RT of unclicked 5’-AzScBc DNA and TSO, as described before. Cas9-gRNA complex (6.6 uM S. pyogenes Cas9 Nuclease (NEB, M0386T), 20 uM gRNA, 1x NEBuffer r3.1, and Nuclease-Free Duplex Buffer (IDT, 11-05-01-04)) was prepared by incubation for 15 min at 25°C. The incubated complex was added to the cleaned PCR reaction and incubated for 1 hour at 37°C.

### PCR amplification and NGS

The DASHed library was PCR amplified with NEBNext Ultra II Q5 High-Fidelity 2X Master Mix (NEB, M0544) and 0.5 uM PCR primers with unique dual index using the two-step PCR cycles:

|  | Steps | Denature<br>(98°C) | Anneal<br>(60°C) | Extension<br>(72°C) | Storage<br>(4°C) |
| --- | --- | --- | --- | --- | --- |
|  | Initial denaturation | 30 sec | - | - | - |
| 6 cycles | Denaturation | 10 sec | - | - | - |
|  | Anneal |  | 20 sec | - | - |
|  | Extension |  |  | 30 sec |  |
|  | Final extension | - | - | 2 min | - |
|  | Storage | - | - | - | Forever |

The NGS library was sequenced on Illumina NovaSeq SP100 flow-cell with 64 nucleotides Forward Read, 43 nucleotides Reverse Read, 8 nucleotides Index 1, and 8 nucleotides Index 2.

### Alignment & pre-processing

Adaptor sequences were removed from paired-end fastq files using Cutadapt^8^. Precisely, the Read 1 sequence CCCCTGTCTCTTATACACAT and the Read 2 sequence AGATCGGAAGAGCGTCGTGT were trimmed with a maximum error rate of 0.15, requiring a minimum overlap of 12 nucleotides between the read and adapter. The resulting adapter-trimmed reads demultiplexed using Flexbar^9^. Cell barcodes and Unique Molecular Identifiers (UMIs) were extracted from the 5’ end of Read 1, applying a barcode error rate of 0.15 and retaining reads of at least 14 nucleotides in length. The adapter-clipped and demultiplexed reads were first mapped to the mouse ribosomal genome using bowtie2^10^ in --very-sensitive mode. The reads unmapped to the ribosomal genome were mapped to the mouse genome (mm10 build) in --very-sensitive mode. After mapping, duplicate reads were identified and removed utilizing UMI and mapping coordinates with UMI-tools^11^.

### Estimation of capture efficiency

The average capture efficiency of scGRO-seq was estimated to be approximately 10% by two methods. First, we used data in the intron seqFISH study^12^, which quantified the abundance of 34 introns by single-molecule fluorescent in-situ hybridization (smFISH). Based on the slope of the line of best fit between data from smFISH and intron seqFISH, the detection efficiency of intron seqFISH was estimated to be 44%. When scGRO-seq was compared with intron seqFISH, the detection efficiency of scGRO-seq was 23% of intron seqFISH. Based on these two detection efficiencies, the estimated capture efficiency of scGRO-seq is ∼10% (23% of 44% is approximately 10%).

For a second independent estimate, we used a set of studies that quantified the number of actively transcribing RNA polymerases^13,14^. In these experiments, cells trapped in agarose were permeabilized, RNA was digested, and the RNA polymerases were allowed to incorporate ^32^P uridine triphosphate. The quantification of total ^32^P uridine incorporation and the determination of the average length of extended transcripts indicated that the average number of RNA polymerase II in Hela (human cervical cancer) cells is ∼65,000. We found that 20% of the transcriptionally engaged RNA polymerase II are present in a paused state around the promoter-proximal and polyadenylation sites, which are undetected by nuclear run-on reaction in the absence of a high concentration of ionic detergent^4^, such as scGRO-seq. Excluding the paused regions, scGRO-seq captured 1,735 transcripts on average per cell in the gene-bodies. Further accounting for the 14% bigger human genome and, more importantly, a mean 2.2-fold increase in transcription in cancer cells^15^, the estimated number of discoverable transcribing RNA polymerase II molecules in gene bodies of a mouse cell is in the range of ∼20,000. 1,735 transcripts per cell results in ∼10% capture efficiency.

### Enhancer annotation

Active transcription regulatory elements (TREs) in mESCs were identified with PRO-seq data using dREG^16^. Further filtering of the dREG carried out to eliminate TREs within or proximal to 1500 bp of the RefSeq annotated genes (n = 23,980) identified 68,299 high-confidence TREs. The remaining TREs within 500 bp of each other were combined, resulting in the final list of 12,542 enhancers. In order to capture nascent RNA derived from elongating RNA polymerases at these enhancers, the TREs were extended at least 1500 bp from the transcription start site (TSS) in both directions. The overlapping enhancers after extension were stitched together.

### Evidence of bursting

Transcriptional bursting was examined *de novo* using scGRO-seq data by measuring two parameters - the multiplicity of RNA polymerases and the distance between the RNA polymerases. The bursting model suggests that transcription occurs in short bursts punctuated by long silent periods, resulting in “on” and “off” states. The alternative model is the relatively uniform transcription initiation by primarily solitary RNA polymerase. We expected two observations under the bursting model.

First, we expected a higher incidence of more than one RNA polymerase per burst and a concurrent depletion of single RNA polymerases. To test the evidence of bursting, we selected genes longer than 11 kb (n = 13,564) and trimmed 0.5 kb regions from the gene’s 5’ and 3’ ends that are known to harbor paused polymerases. With an average transcription rate of 2.5 kb/min, the remaining 10 kb region resulted in an observation window of four minutes. Based on the evidence of monoallelic transcription described in the main text and a short observation window of four minutes, we assigned all signals for a gene in individual cells to one allele. We quantified the observed incidence of zero, one (singlets), and more than one RNA polymerase (multiplets) per allele. The vast majority of alleles had zero polymerase. To calculate the expected incidences of RNA Polymerases under the non-bursting model, we permuted the cell identity of scGRO-seq reads 200 times without changing the read positions. The permutation breaks the bursting-mediated association between RNA polymerases and mimics the RNA polymerases distribution under the non-bursting model. We quantified the permuted incidences of zero, singlets, and multiplets.

Second, if more than one RNA polymerase is observed in the burst window, either due to transcriptional bursting or random chance, we expected the transcription bursting model would result in more closely spaced molecules than expected by the random chance. We took all multiplets in observed or permuted data and calculated the distance between RNA polymerase molecules within each pair. We binned the distances in 50 bp bins and calculated the ratio of RNA polymerase pairs between the observed and permuted data.

### Burst kinetics

Genes over 11 kb (n = 13,564) were selected for studying transcriptional bursting kinetics, and 500 nt regions at both ends known to harbor paused polymerases were truncated. In cases where genes exceeded 10 kb after trimming, they were shortened to 10 kb starting from the gene’s initiation site. With an average transcription rate of 2.5 kb/min, this 10 kb burst window served an average burst duration of 4 minutes. The calculation of burst size and burst frequency proceeded as follows:

Burst Size: For each gene, the number of cells with at least one read within the 10 kb burst window (number of bursts) was identified, and then the average reads per burst was computed. If a consistent single read per burst was observed, that gene’s burst size was set to 1. However, if the average burst size was 1.2, the residual burst above 1 indicated a higher burst size. Accounting for the 10% capture efficiency, where the likelihood of capturing paired reads within a burst window is 1%, the residual burst was proportionally adjusted by the capture efficiency.

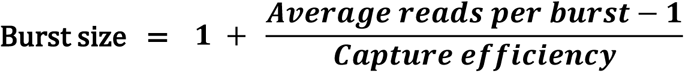

Burst Frequency: For each gene, burst frequency was determined as the number of bursts per allele (two alleles in autosomal and one in sex chromosomes) per transcription time. The transcription time was calculated as the duration needed to traverse the 10 kb burst window with a uniform transcription rate of 2.5 kb/min, translating to four minutes. The calculated burst frequency was normalized by capture efficiency, taking the burst size into account. While burst events with a larger burst size like ten would be consistently detected even with 10% capture efficiency, normalization was applied for cases where a burst size like four would result in a 60% false negative rate, indicating a non-existent burst despite active bursting. Thus, burst frequency normalization was scaled by burst size to ensure accurate quantification.

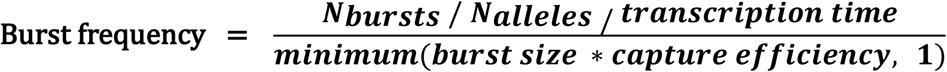

Genes with core promoter elements like TATA and Initiator sequences were retrieved from http://epd.vital-it.ch^17^. Genes containing a pause button, a sequence associated with promoter-proximal paused RNA polymerase, were recovered from the CoPRO dataset^18^.

### Simulation of idealized burst kinetics

We simulated read counts for populations of single cells to evaluate the performance of our estimators for burst rate and size. In the first simulation, we randomly generated the true burst size (T_size_) for all human genes from a normal distribution (mean = 2, standard deviation = 3). Similarly, we generated true burst rates (T_rate_) for all human genes from a normal distribution (mean = 1, standard deviation = 1). T_size_ less than 1 was corrected to 1, and T_rate_ less than 0.1 burst/hour was corrected to 0.1. These parameters were used to simulate reads per gene per cell as follows:

1. For each cell and each gene, sample from a Poisson distribution with rate parameter λ = T_rate_.
2. Scale the sampled burst by T_size_ and round to the nearest integer.
3. After generating molecule counts for all genes and all cells, randomly subsample to a specified level (e.g., 10% sampling efficiency) without replacement.

In the second simulation, T_size_ and T_rate_ were taken from our genome-wide estimates described in Fig. 2, and reads per gene per cell were generated similarly. Simulations were performed ten times to ensure consistent results.

### Cell-cycle analysis

Three sets of transcriptionally characterized genes were used to characterize the cell-cycle phase in individual cells. Transcription of 68 replication-dependent histone genes^19^ on chr3, chr6, chr11, and chr13 were used to determine the S phase collectively. Transcription of four genes (Orc1, Ccne1, Ccne2, and Mcm6) were used to assign G1/S phase and six genes (Wee1, Cdk1, Ccnf, Nusap1, Aurka, Ccna2) were used to assign G2/M phase^20^. Cells with more than a read in one of the genes or reads in more than one gene were hierarchically clustered, which revealed three major clusters of the cell-cycle phase-specific transcription pattern. The other three smaller clusters without distinct transcription patterns were not considered for downstream analyses.

### Gene-Gene co-transcription

The co-transcription of genes was determined by two criteria: correlation and permutation. scGRO-seq reads were collected from up to the first 10 kb of genes after 500 bp regions at both ends were trimmed (n = 15,666). The genes by cells expression matrix was binarized. For the correlation approach, pairwise correlation was performed for all gene pairs, and the p-value was adjusted for multiple hypothesis correction using the Benjamini-Hochberg (BH) method.

Permutation was performed by shuffling the cell IDs of reads while maintaining their gene assignments. The permutation method accounts for several unknown and known biases, such as read depth per cell. The observed and permuted co-transcription frequencies of gene pairs were calculated. The empirical p-value for a gene pair was determined by counting the incidence of equal or higher co-transcription frequency in 1000 permutations compared to the observed co-transcription frequency.

Gene pairs with correlation coefficients of greater than 0.1 and adjusted p-values of less than 0.05 from the correlation approach, and an empirical p-value of less than 0.05 from the permutation method were considered co-transcribed. A network of pairwise co-transcribed genes was created using the Leiden algorithm, and the modules were selected for gene-ontology analyses using the clusterProfiler R package.

### Enhancer-Gene co-transcription

Enhancer-gene co-transcription was determined following the logic of gene-gene co-transcription, substituting genes on one arm with enhancers. scGRO-seq reads were collected from up to the first 10 kb of genes after 500 bp regions at both ends were trimmed, and from at least a 3 kb region around enhancers (1,500 bp sense and 1,500 bp anti-sense) after a 500 bp region around the TSS was removed to avoid paused polymerases. Strand-specific reads on either side of the enhancer TSS were combined to determine enhancer expression. The features (genes + enhancers) by cell expression matrix was binarized, and the co-transcribed enhancer-gene pairs were determined using the correlation and permutation methods, similar to the approach used in the gene-gene co-transcription calculation. Enhancer-gene pairs only from the same chromosomes were retained for downstream analyses. We also included non-overlapping super-enhancers identified in mESCs^21^.

### Enhancers of pluripotency factors

Validated enhancers associated with pluripotency transcription factors Oct4 (Pou5f1), Sox2, Nanog, and Klf4 were collected from numerous studies^22–26^. To define time bins within genes, genes were divided into 5 kb bins (two minutes bins calculated using the 2.5 kb/min constant transcription rate of elongating RNA polymerases) in the sense and antisense direction until the end of the transcription wave called by groHMM^27^, or they overlapped bins from other genes. For enhancers, TSS was first determined based on the strongest Oct4, Sox2, and Nanog chromatin immunoprecipitation and sequencing (ChIP-seq) peaks. The precise position was determined by evaluating the divergent transcription around them. The reads from corresponding bins in sense and antisense directions were combined.

### CRISPR-validates SEs

A set of validated SEs and their target genes were used from a previously published study^28^. SEs in gene introns or associated with miRNA were excluded due to the ambiguity in assigning reads and short gene length, respectively. For the time bin analyses, genes and SEs were divided into four 5 kb bins (two minutes with the 2.5 kb/min constant transcription rate of elongating polymerases) in the sense and antisense direction, limiting the analyses to the first 20 kb. Using a 20 kb region in this analysis yields four 5 kb bins. TSS was first determined based on the strongest Oct4, Sox2, and Nanog chromatin immunoprecipitation and sequencing peaks, and precise position was determined by evaluating the divergent transcription around them. The reads from corresponding bins in sense and antisense directions were combined. The scrambled random pairs in SE-gene time bin analysis represent the co-transcribed bins between SEs and genes that are not the verified pairs.

### External data

Various data types were analyzed, compared, and benchmarked against this study. PRO-seq libraries from (GEO: GSE169044) were prepared with the same cells used for scGRO-seq under identical conditions^29^. Intron seqFISH data on mESCs was downloaded from Table S1 of a published study^12^. The genes by cells matrix was binarized, and burst frequency was calculated assuming the signal in each gene comes from a burst equivalent to the 10 kb region used in scGRO-seq, given the probes were designed against the introns at the 5′ regions of genes. mESCs scRNA-seq was used from a previous study^30^ and the burst kinetics was downloaded from 41586_2018_836_MOESM5_ESM.xlsx file associated with this study.

## Data availability

Sequencing files for scGRO-seq, inAGTuC, and AGTuC experiments are deposited in NCBI’s Gene Expression Omnibus and are accessible through GEO Series accession number GEO: GSE242176.

## Code availability

The codes used in this study are available at https://github.com/jaymahat/scGROseq.

## Acknowledgments

We are grateful to the current and past members of the Sharp laboratory, especially to Dr. Amanda Whipple, for the discussion and critical review of the manuscript. We thank Arjun Bhutkar from the Koch Institute for Integrative Cancer Research for his guidance in computational analyses, Suman Bose for his help with FACS, and Diogo Ribeiro from the University of Lausanne, Switzerland and James Weber from the Broad Institute of Harvard and MIT for their insight on statistical tests on co-transcriptional measurement. We thank the Swanson Biotechnology Center Flow Cytometry Facility for help with FACS, and Stuart Levine and the staff at BioMicro Center at the Koch Institute for Integrative Cancer Research for their help with NGS. This work was supported by Program Project Grant P01-CA042063 from the NCI (P.A.S.) and by United States Public Health Service grants R01-GM034277 from the NIH (P.A.S.). D.B.M. is supported by the Emerald Foundation Postdoctoral Transition Award. He was previously supported by the Gertrude B. Elion Research Fellowship from GSK.

## Authors contributions

D.B.M. and P.A.S. conceived the study. D.B.M., S.K.W., and J.F. optimized click-chemistry and library preparation methods, and prepared NGS libraries. D.B.M. and N.D.T. analyzed the data with the help of S.B. on pre-processing and J.D.M.R. on scRNA-seq and kinetic analyses. D.B.M. and P.A.S. wrote the manuscript, and all co-authors provided feedback. P.A.S. supervised the project.

---

**Extended Data Fig. 1.**
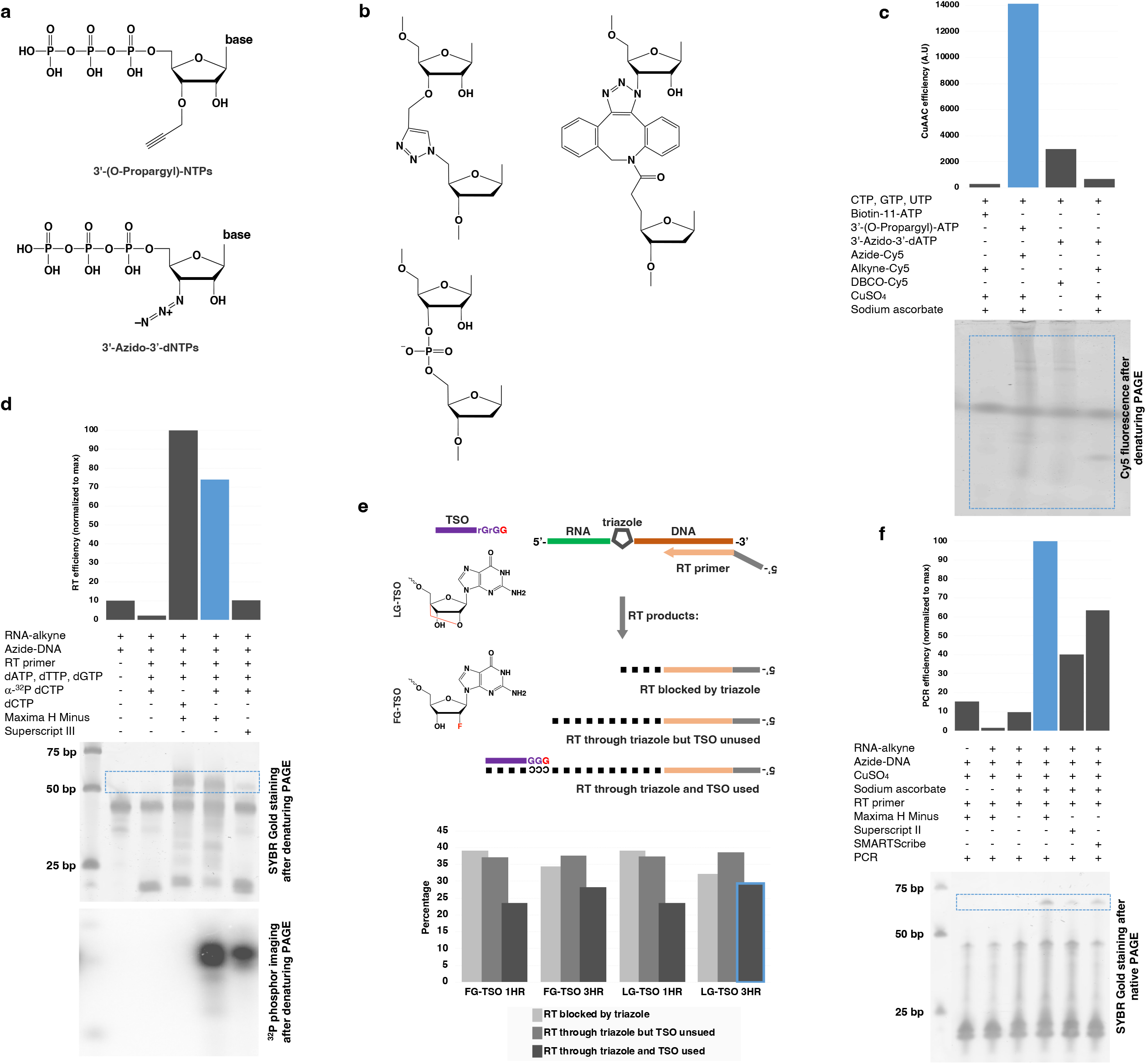
click-chemistry mediated nascent RNA conjugation to single-stranded DNA and optimization of reverse transcription. **a**, Click-chemistry compatible nucleotides tested in AGTuC development. A few nucleotide triphosphates were custom synthesized or sourced with few properties in mind - smaller size, chain termination ability, and the possibility of incorporation by native RNA polymerases. **b,** Structure of the triazole linkage formed by CuAAC between the nascent-RNA terminally labeled with 3’-(O-Propargyl)-NTPs and the azide-labeled DNA (top left), the linkage formed by SPAAC between the nascent-RNA terminally labeled with 3’-Azido-3’-dNTPs and DBCO DNA (right). The phosphodiester linkage in a native oligonucleotide is shown for comparison (bottom left). **c,** Incorporation efficiency of 3’-(O-Propargyl)-ATP or 3’- Azido-3’-dATP by native RNA polymerase in nuclear run-on reaction. The propargyl or azide labeled nascent RNA is clicked with Cy5 via CuAAC (Azide-Cy5 or Alkyne-Cy5) or SPAAC (DBCO-Cy5), resolved in a denaturing polyacrylamide gel electrophoresis (PAGE), and quantified by measuring the Cy5 fluorescent from the gel image. The blue dotted line represents the gel region that was quantified. **d,** Relative quantification of reverse transcription (RT) efficiency of two commercial enzymes traversing through the triazole link formed between the alkyne-labeled RNA and azide-labeled DNA by CuAAC. RT was performed in the presence of either native dCTP or radioisotope a-^32^P dCTP, and the RT reaction was resolved in denaturing PAGE and imaged sequentially for nucleic acid signal (top gel) and radioisotope signal (bottom gel). **e,** Quantification of aborted intermediate and completed desired products (RT through triazole and TSO used) formed during the one hour or three hours of RT using TSO with terminal Locked-Nucleic-Acid-Guanosine (LG) or 2’-Fluoro-Guanosine (FG). **f,** Confirmation and relative quantification of CuAAC, RT, and PCR of clicked product formed between the alkyne-labeled RNA and azide-labeled DNA by three commercial Reverse transcriptase enzymes. **Note:** The blue bar, line, or border represents the “winner” condition.

**Extended Data Fig. 2.**
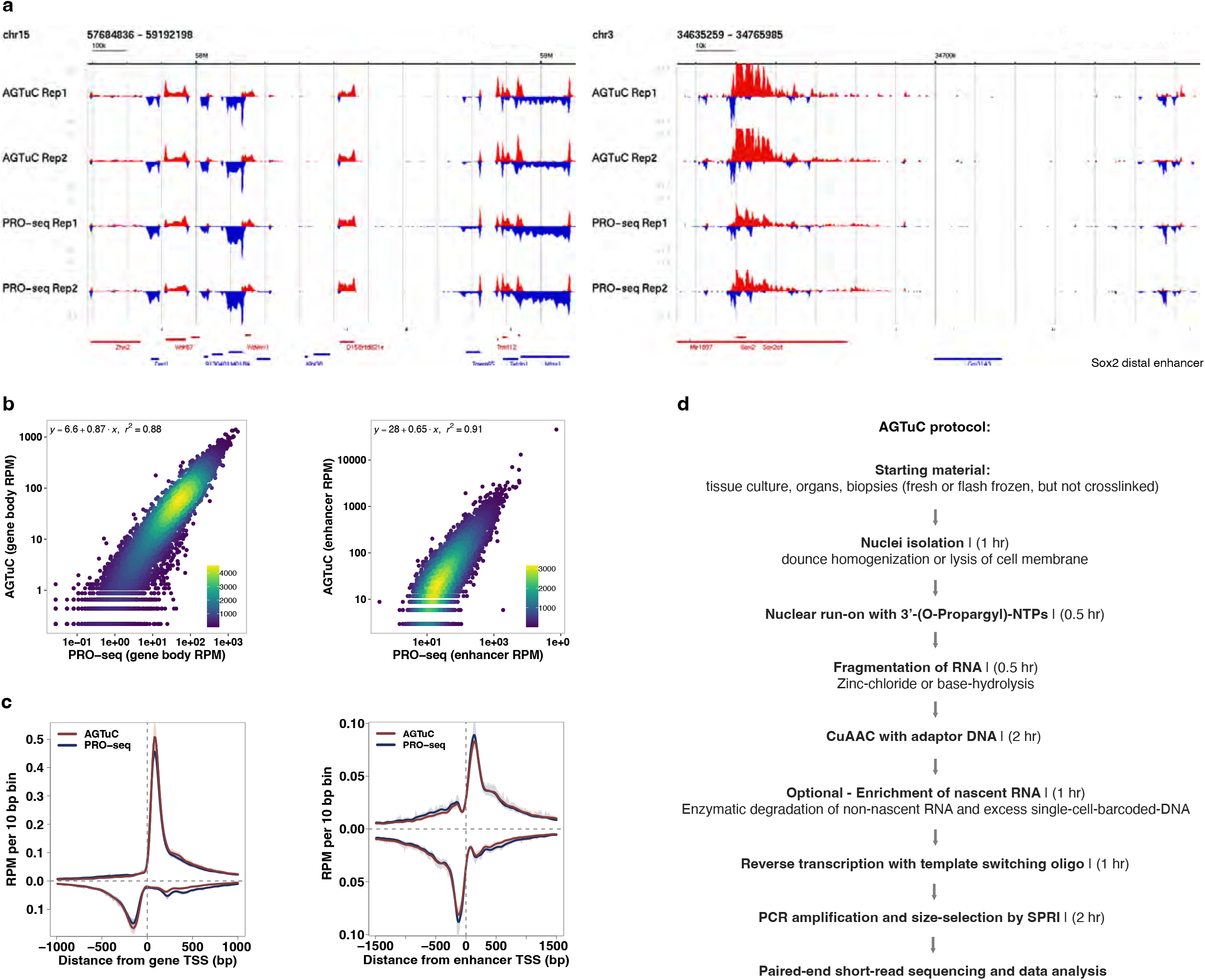
Comparison between AGTuC and PRO-seq. **a**, Representative genome-browser screenshots with two replicates of AGTuC and PRO-seq showing a region in chromosome 15 (left) and a region in chromosome 3 containing the Sox2 gene and its distal enhancer (right) of the mouse genome (mm10). **b**, Correlation between AGTuC and PRO-seq reads per million sequences in gene bodies (left, n = 19,961) and enhancers (right, n = 12,542). **c**, Metagene profiles of AGTuC and PRO-seq reads per million per 10 base pair bins around the TSS of genes (left, n = 19,961) and enhancers (right, n = 12,542). The line represents the mean, and the shaded region represents 95% confidence interval. **d,** Major steps with the approximate time required in AGTuC library preparation.

**Extended Data Fig. 3.**
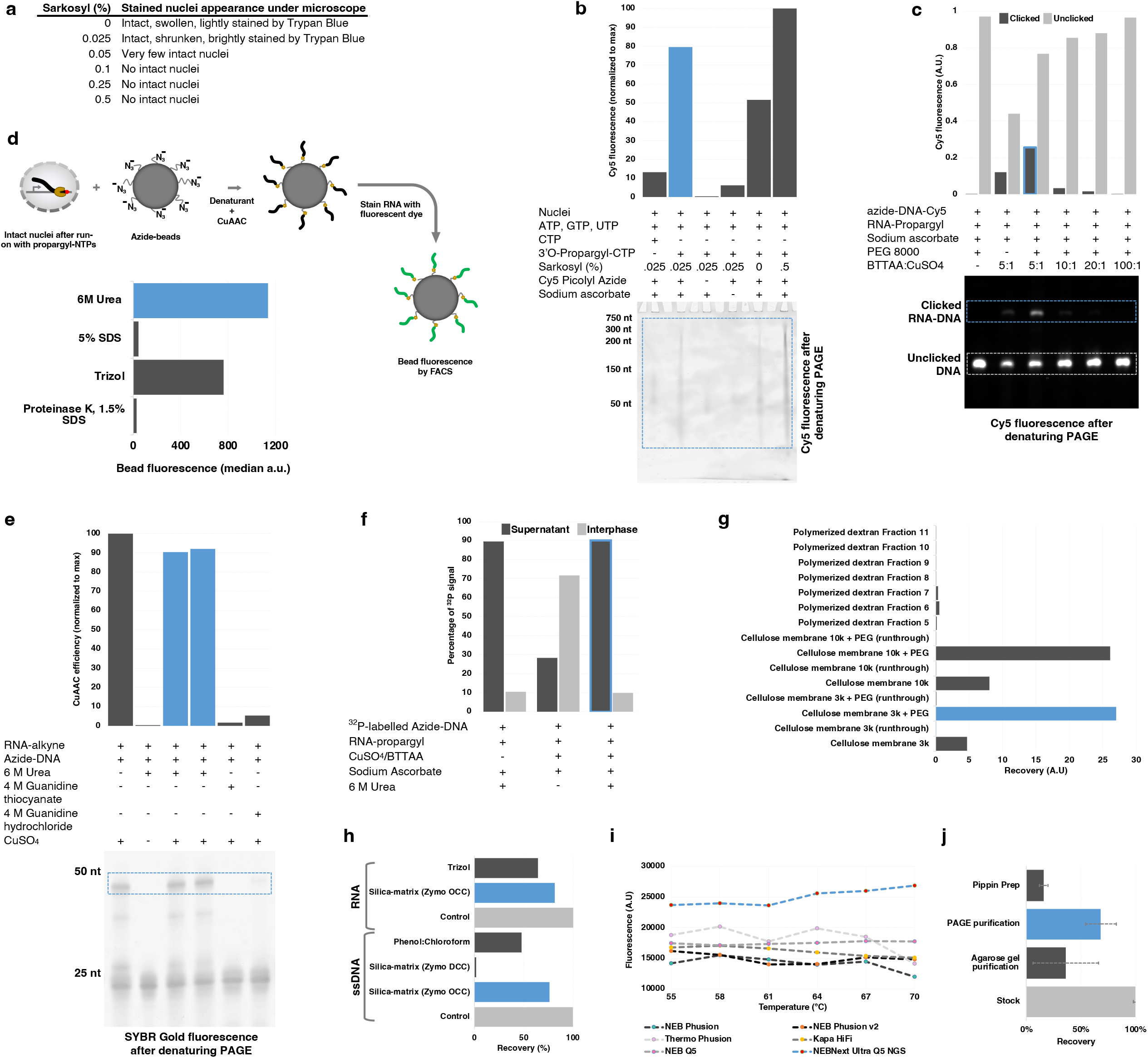
Optimization of intact nuclei run-on reaction and NGS library preparation steps. **a**, Physical appearance of Trypan Blue stained nuclei under microscope treated with various sarkosyl concentrations. **b,** Relative quantification of nuclear run-on efficiency with various sarkosyl concentrations. Nascent RNA collected after nuclear run-on reaction with either native CTP or click-compatible 3’O-Propargyl- CTP was clicked with Cy5-azide, resolved in denaturing PAGE, and images for Cy5 fluorescence. **c,** Effect of different ratios of CuAAC accelerating ligand BTTAA in CuAAC efficiency. RNA-propargyl was clicked with azide-DNA containing Cy5 in the presence of various ratios of BTTAA:CuSO_4_, resolved in denaturing PAGE, and images for Cy5 fluorescence. **d,** Relative quantification of denaturing efficiency of commonly used denaturing agents to release the nascent RNA from RNA polymerase complex. Intact nuclei after run-on with 3’O-Propargyl-NTPs were treated with denaturing agents in the presence of azide-labeled beads and CuAAC reagents, allowing nascent RNA to click with the beads. Beads were stained with RNA-binding dye and measured for fluorescence by FACS. **e,** Effect of denaturing agent’s presence in CuAAC efficiency. The blue outline in the image of denaturing PAGE denotes the click product between the RNA-alkyne and azide-DNA. **f,** Role of urea in the residence of clicked RNA-DNA conjugate in either supernatant or interphase of Trizol during the clean-up of CuAAC reaction, as quantified by the scintillation count of ^32^P radioisotope. **g,** Desalting (removal of CuSO4, BTTAA, and sodium ascorbate from CuAAC reaction) efficiency of polymerized dextran and cellulose membrane. Fluorescence from Cy5-labeled RNA-DNA conjugate was measured in elution fractions from columns packed with polymerized dextran and elution from different pore-size cellulose membrane centrifugation tubes with or without PEG 8000. **h,** Relative recovery of ssDNA or RNA from phenol:chloroform or silica-based matrix column purification. Clicked RNA-DNA conjugate was radioisotope labeled using Polynucleotide kinase and γ-^32^P ATP, and the cleaned reaction was quantified using a scintillation counter. **i,** PCR amplification efficiency of clicked RNA-DNA conjugate using different commercial PCR amplification kits. The PCR reaction was resolved in native PAGE, stained with SYBR Gold, and quantified using ImageJ software. **j,** Relative recovery of size-selected dsDNA. A mock NGS library (purified PCR product) was selected for the desired size using various size-selection methods, and the recovered dsDNA was quantified using a dsDNA-specific fluorescence kit (Qubit). **Note:** The blue bar, line, or border represents the “winner” condition.

**Extended Data Fig. 4.**
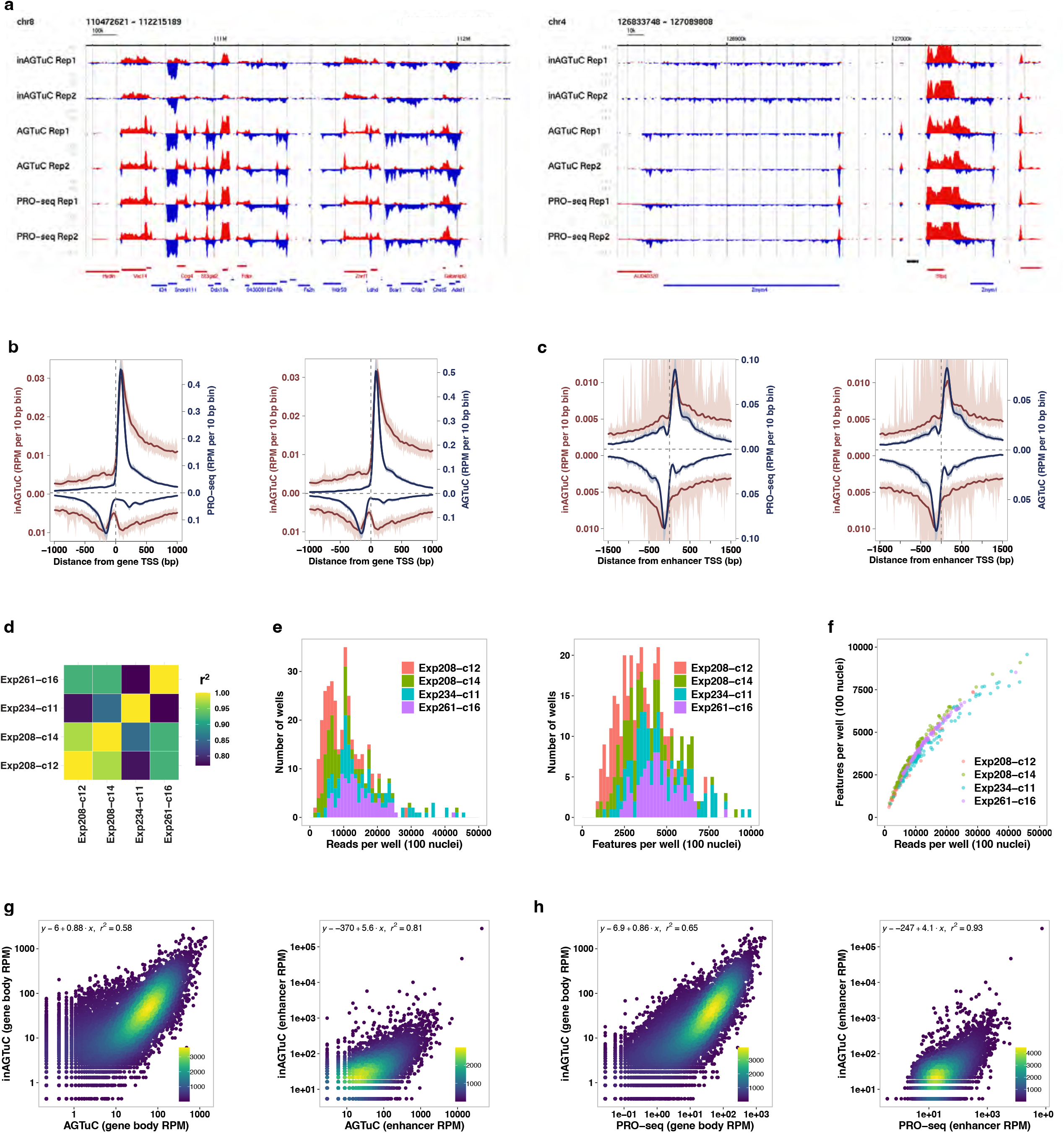
Benchmarking inAGTuc against AGTuC and PRO-seq. **a**, Representative genome-browser screenshots with two replicates of inAGTuC, AGTuC, and PRO-seq showing a region in chromosome 8 (left) and a region in chromosome 4 of the mouse genome (mm10). **b, c,** Comparison of inAGTuC metagene profiles with PRO-seq and AGTuC using reads per million per 10 base pair bins around **(b)** the TSS of genes (n = 19,961) and **(c)** enhancers (n = 12,542). The line represents the mean, and the shaded region represents 95% confidence interval. **d**, Correlations of inAGTuC reads per million sequences in gene bodies (n = 19,961) between the four replicates. **e**, Distribution of reads per well (left) and features per well (right) in four replicates of 96-well plate inAGTuC libraries. Each well contains 100 nuclei. **f**, Relationship between the reads per well and the number of features detected per well in four replicates of 96-well plate inAGTuC libraries. **g, h**, Correlation between inAGTuC and AGTuC (two left panels) or PRO-seq (two right panels) reads per million sequences in **(g)** the body of genes (n = 19,961) and **(h)** enhancers (n = 12,542).

**Extended Data Fig. 5.**
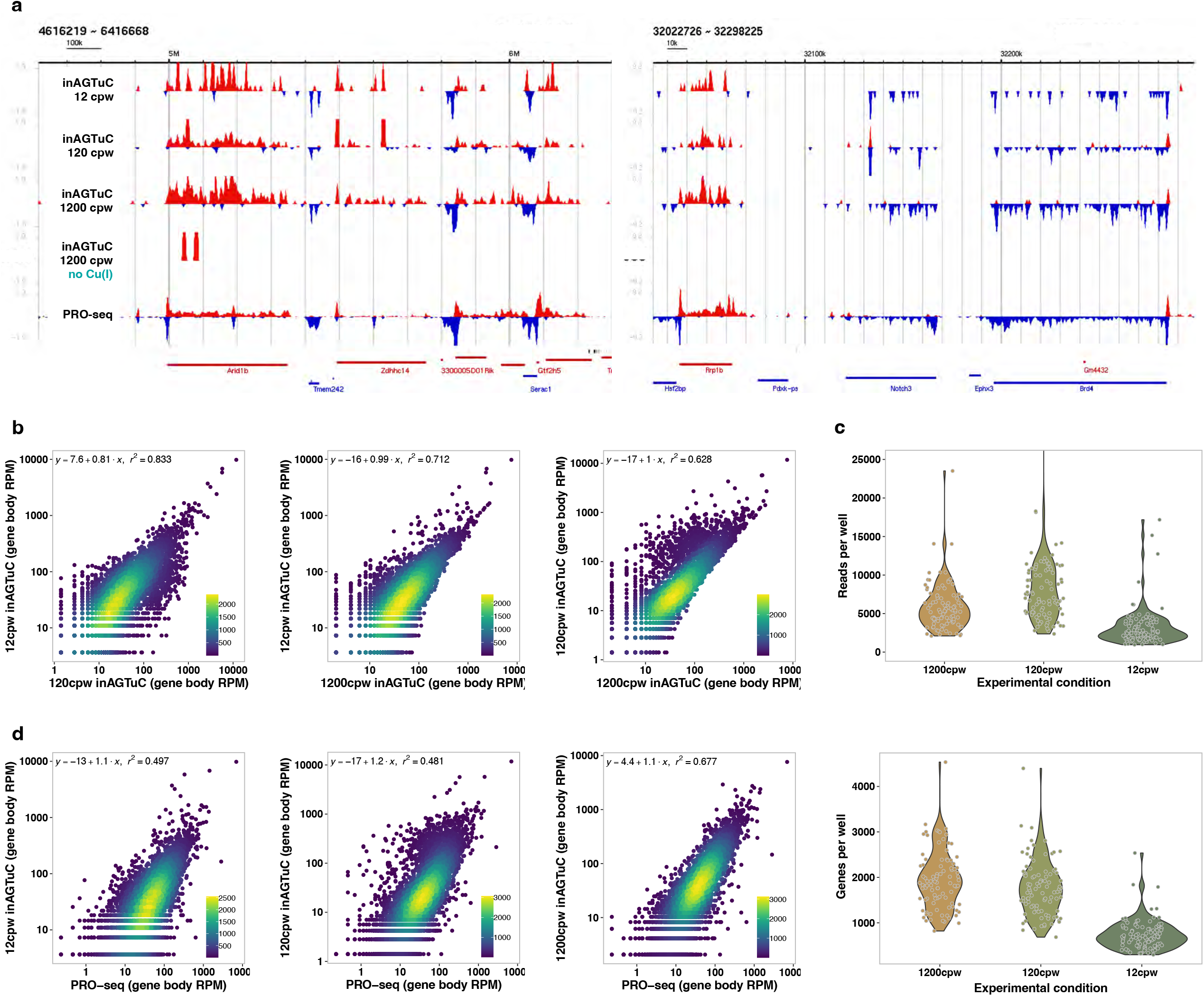
Feasibility demonstration of inAGTuC with fewer cells. **a**, Representative genome-browser screenshots of inAGTuC library from one 96-well plate with each well containing either 12, 120, or 1200 cells per well (cpw) showing two regions in chromosome 17 of the mouse genome (mm10). inAGTuC library with 1200 cpw but without Cu(I) in the CuAAC reaction (fourth track) and the PRO-seq library (fifth track) serve as the negative and positive control, respectively. **b**, Correlations among 12 cpw, 120 cpw, and 1200 cpw inAGTuC libraries in the body of genes (n = 19,961). **c**, Distribution of reads per well (top) and genes per well (bottom) in 12 cpw, 120 cpw, and 1200 cpw inAGTuC libraries. **d**, Correlations between PRO-seq and 12 cpw, 120 cpw, and 1200 cpw inAGTuC libraries in the body of genes (n = 19,961).

**Extended Data Fig. 6.**
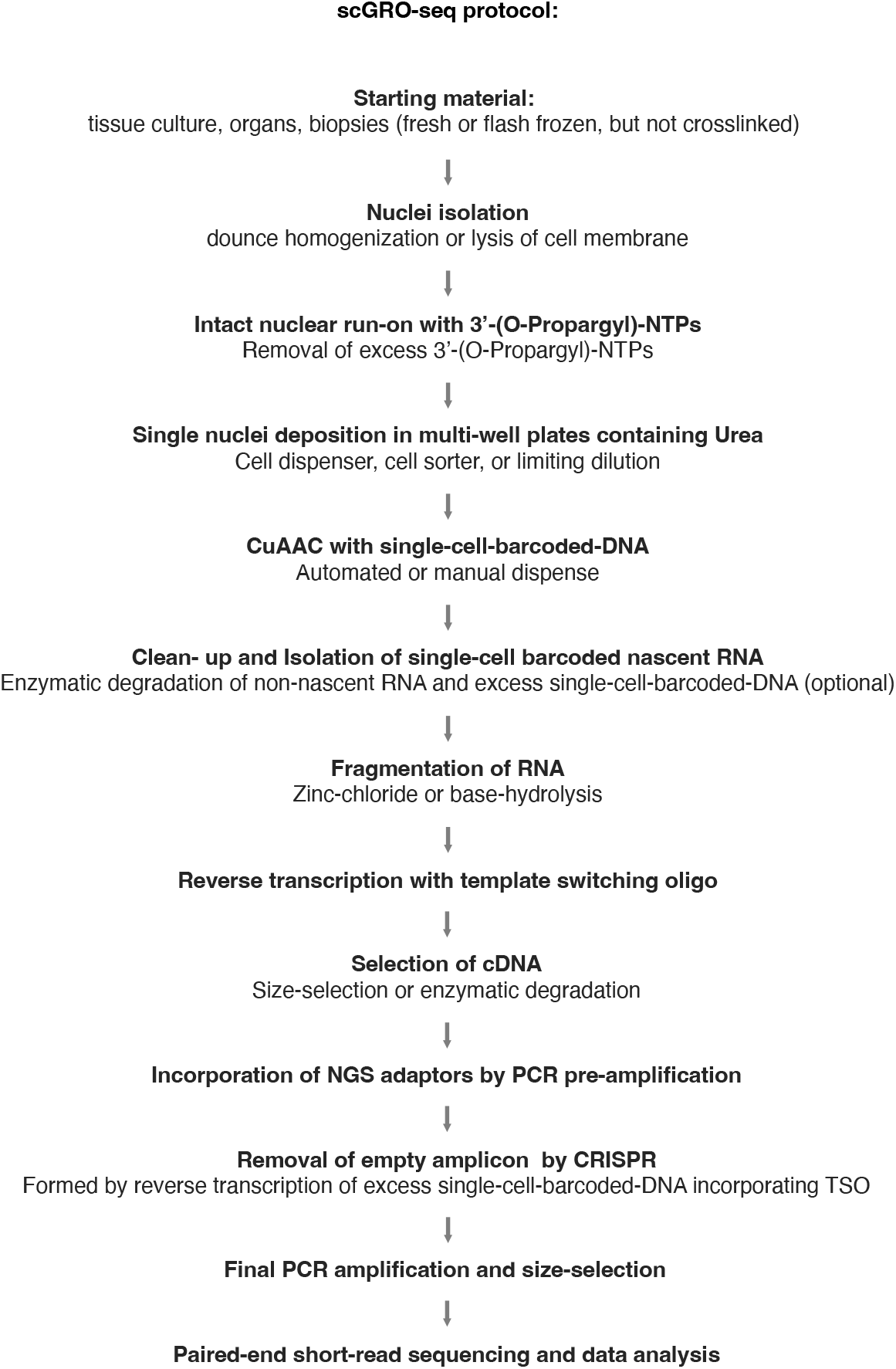
scGRO-seq library preparation. Major steps involved in scGRO-seq library preparation.

**Extended Data Fig. 7.**
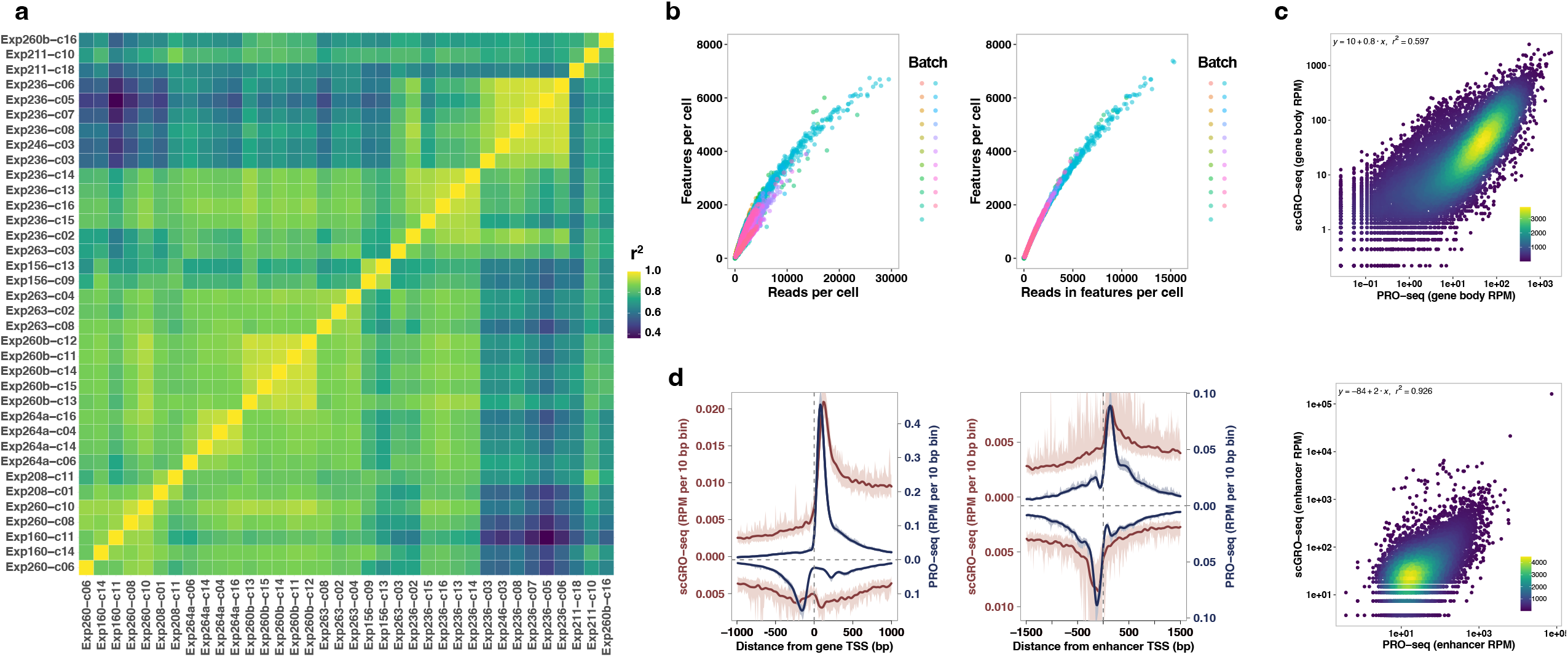
Benchmarking scGRO-seq. **a**, Coefficient of determination (r^2^) between each 96-well plate from scGRO-seq batches that passed the quality-control threshold. r^2^ was calculated from average reads per 96 cells in all genes and enhancers. **b**, Relationship between the number of features detected per cell and the scGRO-seq reads per cell (left), or scGRO-seq reads in features per cell (right). **c**, Correlation between scGRO-seq and PRO-seq reads per million sequences in gene bodies (top, n = 19,961) and enhancers (bottom, n = 12,542). **d**, Comparison of metagene profiles between scGRO-seq and PRO-seq reads per million per 10 base pair bins around the TSS of genes (left, n = 19,961) and enhancers (right, n = 12,542).

**Extended Data Fig. 8.**
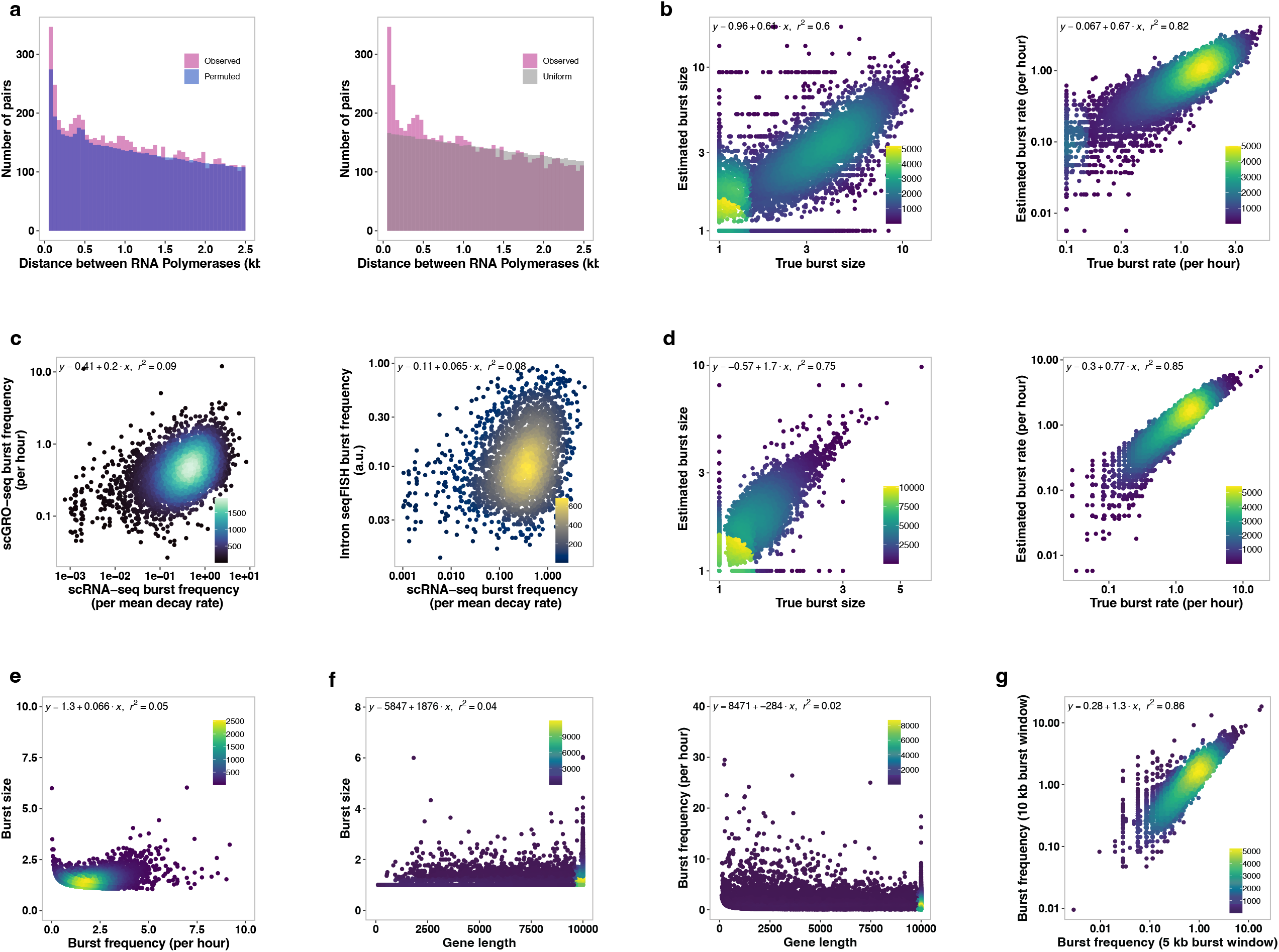
Effect of transcription level, gene length, and burst duration in transcription burst kinetics. **a,** Distribution of distances between consecutive RNA polymerases in the first 10 kb of the gene body in single cells compared with distances from permuted data (randomized cell ID but unchanged read position, left) or uniform data (randomized read position along the gene but unchanged cell ID, right). Distances up to 2.5 kb are shown. **b,** Test of our burst kinetics model by simulating burst size and burst frequency. **c,** Correlation between the burst frequency from scGO-seq (left) and intron seqFISH (right) with the burst frequency from scRNA-seq. **d,** Test of our burst kinetics model by simulation using inferred burst size and burst frequency. **e,** Correlation between the burst size greater than one and the burst frequency of genes. **f,** The effect of gene length (from 100 bp to 10 kb after trimming 500 bp on either end of the genes) on burst size and frequency. **g,** Correlation between burst frequencies calculated from the burst window of either the first 5 kb or the first 10 kb gene bodies.

**Extended Data Fig. 9.**
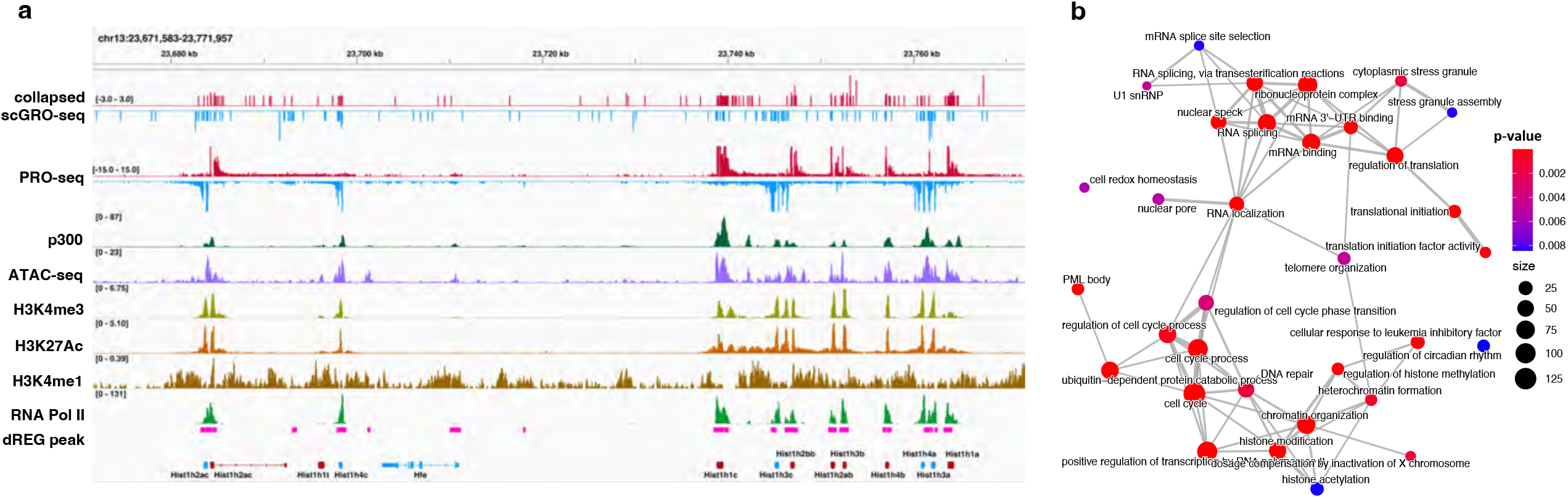
Co-transcription of genes with shared promoters. **a,** Genome-browser screenshot of the histone locus body in mouse chromosome 13 showing transcription of replication-dependent histone genes. **b,** Network of enriched gene ontology terms in co-transcribed genes. A connecting gray line represents at least a 10% overlap of genes between the GO terms. The color of the dots represents the p-value, and the dot size represents the number of contributing genes in the GO term.

**Extended Data Fig. 10.**
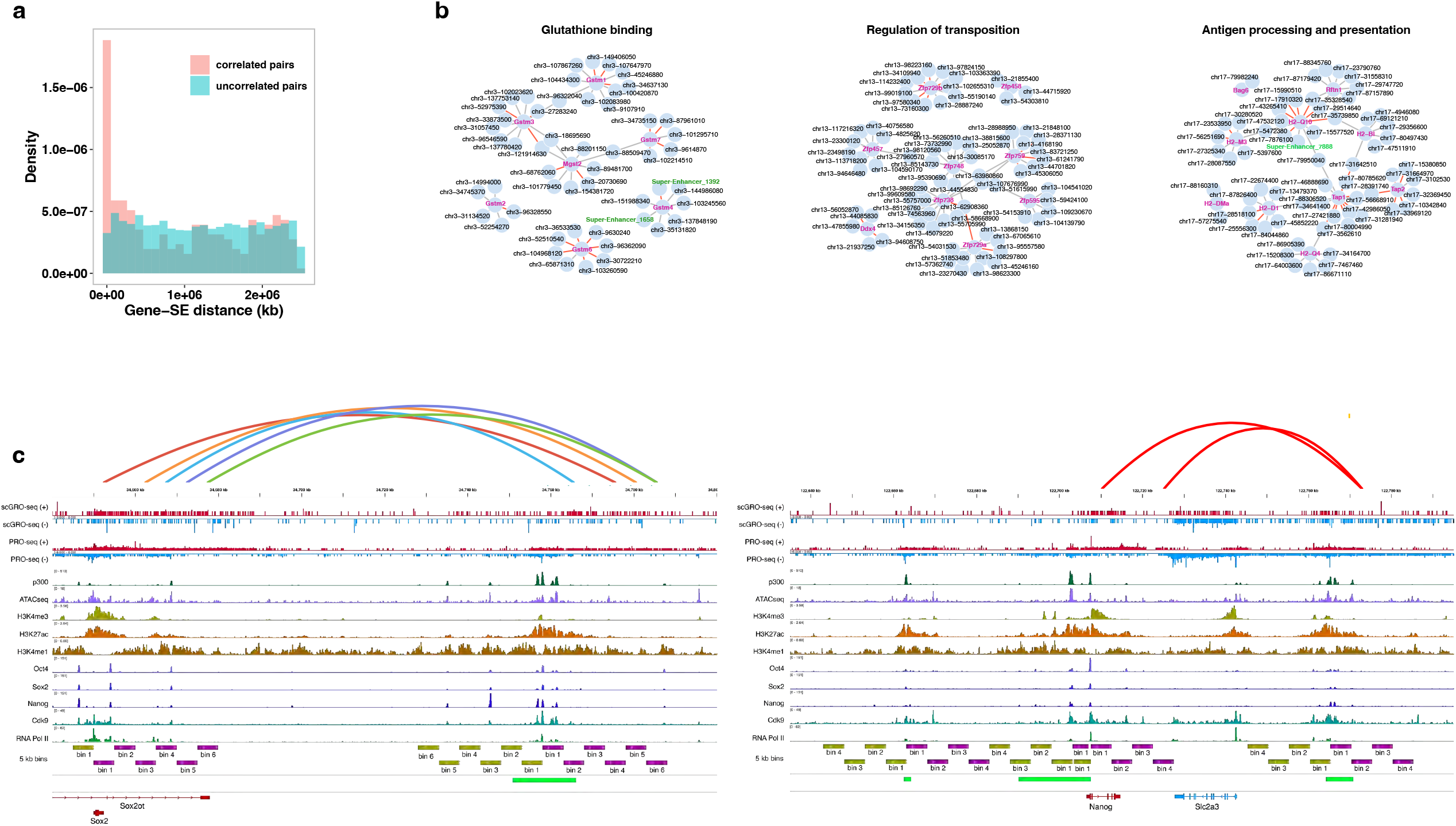
Organization of enhancer-gene co-transcription networks. **a,** Distance between correlated and non-correlated SE-gene pairs within 2.5 Mb of each other. **b,** Co-transcription network of functionally related genes clustered together on the same chromosome shown as examples. Red edges between the enhancer-gene pairs indicate rho > 0.15, and rho > 0.1 and < 0.15 are shown in gray. **c,** Co-transcription between the Sox2 gene and its distal enhancer (left), and the Nanog gene and its three enhancers (right). Green bars represent the annotated SE regions, and the 5 kb bins in sense and antisense strands are represented in magenta and yellow-green bars.

